# Disruption of hippocampal-prefrontal neural dynamics and risky decision-making in a mouse model of Alzheimer’s disease

**DOI:** 10.1101/2024.09.17.613376

**Authors:** Eun Joo Kim, Sanggeon Park, Bryan P. Schuessler, Harry Boo, Jeiwon Cho, Jeansok J. Kim

## Abstract

This study investigates how amyloid pathology influences hippocampal-prefrontal neural dynamics and decision-making in Alzheimer’s disease (AD) using 5XFAD mice, a well-established model system characterized by pronounced early amyloid pathology. Utilizing ecologically-relevant “approach food-avoid predator” foraging tasks, we observed that 5XFAD mice exhibited persistent risk-taking behaviors and reduced adaptability to changing threat conditions, indicative of impaired decision-making. Multi-regional neural recordings revealed rigid hippocampal CA1 place-cell fields, decreased sharp-wave ripple (SWR) frequencies, and disrupted medial prefrontal-hippocampal connectivity, all of which corresponded with deficits in behavioral flexibility during spatial risk scenarios. These findings highlight the critical role of SWR dynamics and corticolimbic circuit integrity in adaptive decision-making, with implications for understanding cognitive decline in AD in naturalistic contexts. By identifying specific neural disruptions underlying risky decision-making deficits, this work provides insights into the neural basis of cognitive dysfunction in AD and suggests potential targets for therapeutic intervention.

## INTRODUCTION

Alzheimer’s disease (AD) research has traditionally focused on memory impairments, but evidence shows AD also disrupts decision-making under risk and ambiguity,^1–10^ essential for daily life. Notably, decision-making deficits may precede memory loss,^11–20^ with early signs like reduced scam awareness and impaired financial judgment linked to amyloid accumulation and heightened AD risk.^11,19,21^ Early to moderate-stage AD patients show impaired decision-making in neuroeconomic tasks,^20,22^ suggesting it could serve as an early marker for AD. Unlike basic memory processes, decision-making requires integrating spatial context, emotional regulation, and past experiences.^23–25^ Risky decision-making may offer a sensitive measure of early AD changes, but animal studies are limited^26^ due to the lack of naturalistic paradigms.

While structural anomalies like amyloid β (Aβ) plaques and tau tangles are well-documented in AD brains, real-time disruptions in neural circuit activity during behavior remain poorly understood. This gap persists despite the limited success of Aβ-targeted therapies^27–29^ and evidence of cognitive resilience in the presence of amyloid pathology.^30–32^ These findings suggest that effective treatments should address both plaques and neural connectivity.^33–35^

Decision-making relies on medial prefrontal cortex (mPFC) interactions with the dorsal hippocampus (dHPC). Damage or inactivation of these regions impairs risk-based decisions in rodents, monkeys, and humans.^36–41^ fMRI studies in humans and single-unit recordings in animals performing probabilistic/delay-discounting tasks link decision-making to mPFC and dHPC activity.^38,42^ AD patients also show deficits in financial decision-making^22^ and set-shifting tasks,^3^ attributed to Aβ plaques and tau tangles in the dHPC and mPFC—regions particularly vulnerable to AD.

Despite these insights, decision-making within ecologically-relevant paradigms and mPFC-dHPC circuit activity remain unexplored in AD animal models. Our recent work documented neural activity in rats during goal-directed foraging in risky environments.^43,44^ We found that dHPC place cells encoded spatial-danger gradients, with unstable firing near a predatory robot but stable firing near the safe nest, highlighting the dHPC’s role in spatial-danger coding. Concurrently, mPFC neurons tracked food procurement and exhibited heightened activity during cautious approaches to a stationary predator, underscoring the mPFC’s role in threat assessment and decision-making. These findings support the utility of naturalistic foraging paradigms to investigate risky choices^3,5,45^ and mPFC-dHPC activity in AD models.

This study employs the 5XFAD transgenic mouse model, which exhibits amyloid pathology from 4 months due to overexpression of amyloid precursor protein (APP) and presenilin 1 (PS1) mutations.^46–49^ Our focus is on dHPC sharp-wave ripples (SWRs) and mPFC-dHPC connectivity. SWRs are brief, high-frequency (150-250 Hz) oscillations related to decision-making, emotional responses, and memory consolidation,^50,51^ and functionally interact with mPFC activity.^52–54^ SWRs are reduced in APP/PS1,^55^ ApoE4-KI (apolipoprotein E4 knockin),^56,57^ and 5XFAD^58^ models, with recent evidence linking them to hippocampal inhibitory synaptic deficits.^59^

By leveraging naturalistic environments that influence and reshape neural activity and behavior,^60–63^ we adapted the rat ‘approach food-avoid predator’ conflict paradigm, ^64,65^ to the 5XFAD model. This approach enables a comprehensive analysis of how AD affects cognition and mPFC-dHPC neural coordination during risky decision-making. Our study aims to provide insights into the neural dynamics and behavioral patterns associated with risky decision-making in AD.

## RESULTS

### 5XFAD mice exhibit risk-prone foraging in an escalating predatory task

5XFAD and WT mice were tested in a foraging task where predatory risk increased with distance from a safe nest (Figure 1). The task consisted of three phases: nest habituation, baseline foraging, and predator testing. During baseline foraging, both 5XFAD and WT mice left the nest and retrieved food pellets with comparable efficiency (Figure S1A). The next day, animals were exposed to a predator (weasel) challenge (Figures 1A and 1B). Data from different sex and age groups were pooled due to consistent behavioral patterns (Figures S1B and S1C).

**FIGURE 1.**
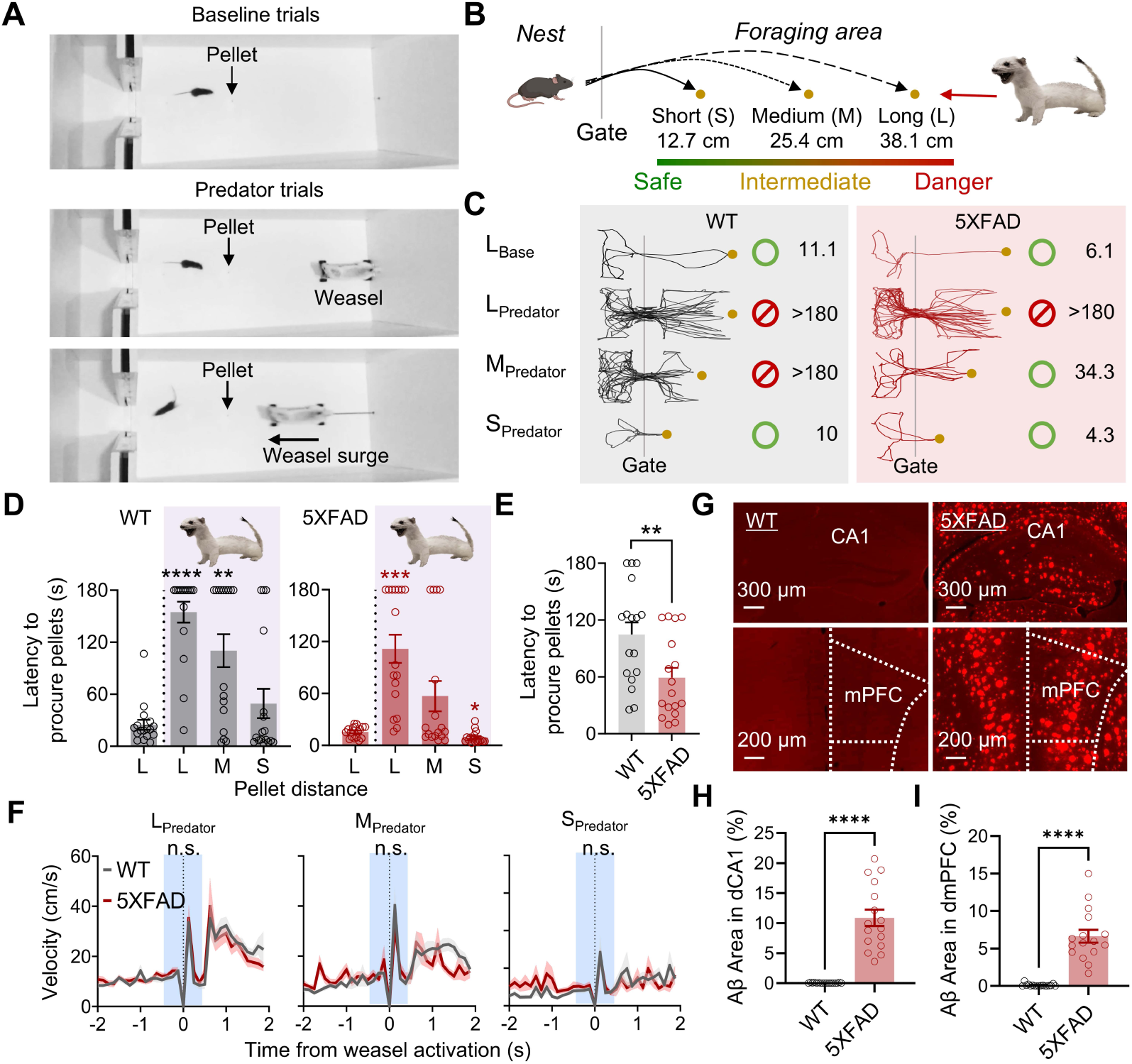
5XFAD mice exhibit risk-prone foraging in an escalating predatory task. (A) Image of a foraging mouse encountering a predatory weasel. (B) Schematic of the behavioral procedure. (C) Representative foraging trajectories for WT and 5XFAD mice during predator testing. (D) Latency to procure pellets in WT (n = 17; *Left*) and 5XFAD (n = 17; *Right*) mice. (E) Mean latency to procure pellets across three weasel trials in WT and 5XFAD mice. (F) Approach (prior to weasel activation) and escape (after weasel activation) velocities in WT and 5XFAD mice at long (L), medium (M), and short (S) distances. No significant group differeneces were observed in approach or escape speeds across the three distance trials (shaded area: −500 to +500 ms relative to weasel activation). (G) Representative images showing Aβ plaque accumulation in the dCA1 and mPFC of a 5XFAD mouse, with no detectable Aβ in a WT mouse. (H,I) Mean area percentage of Aβ accumulation in dCA1 (H) and mPFC (I) of WT and 5XFAD mice. Data are presented as mean ± SEM. \*\**P* < 0.01, \*\*\**P* < 0.001, \*\*\*\**P* < 0.0001.

On the first day of predator testing, both groups initially fled into the nest in response to the surging weasel, showing significantly longer latencies to procure pellets at the 38.1 cm distance compared to baseline (Figures 1C and 1D). At 25.5 cm, WT mice displayed risk-averse behavior, whereas 5XFAD mice showed risk-prone foraging. At the shortest distance (12.7 cm), both groups successfully retrieved the pellet. WT mice demonstrated latencies comparable to their baseline, while 5XFAD mice showed shorter latencies. Across all distances, 5XFAD mice had shorter mean latencies to retrieve pellets compared to WT mice (Figure 1E).

Despite differences in foraging latency, no significant group differences were observed in running speed (velocity), latency to enter foraging area, or the number of attempts at pellets (Figures 1F and S1D-S1F), indicating predator responsiveness and pellet motivation remain unaffected. Histological analysis revealed higher levels of Aβ plaque deposits in the dCA1 and mPFC compared to WT mice (Figures 1G-1I). These findings suggest that Aβ accumulation in 5XFAD mice disrupts their ability to discern safety boundaries under escalating predatory risk, promoting risk-prone foraging.

### 5XFAD mice exhibit inflexible foraging and rigid place cell coding in a conditional predatory task

To assess the impact of Aβ plaques on decision-making and hippocampal spatial coding, 5XFAD and WT mice were implanted with tetrode arrays in the dHPC (targeting the CA1 subregion) and the mPFC ipsilaterally (Figure S2A). Mice underwent successive stages of nest habituation, baseline foraging, and predator testing within a T-shaped maze (Figure 2A) featuring two distinct pellet types (grain-based and chocolate-flavored) located in separate goal arms. Neural recordings were collected during pre-predator and predator sessions (Figure 2B).

**FIGURE 2.**
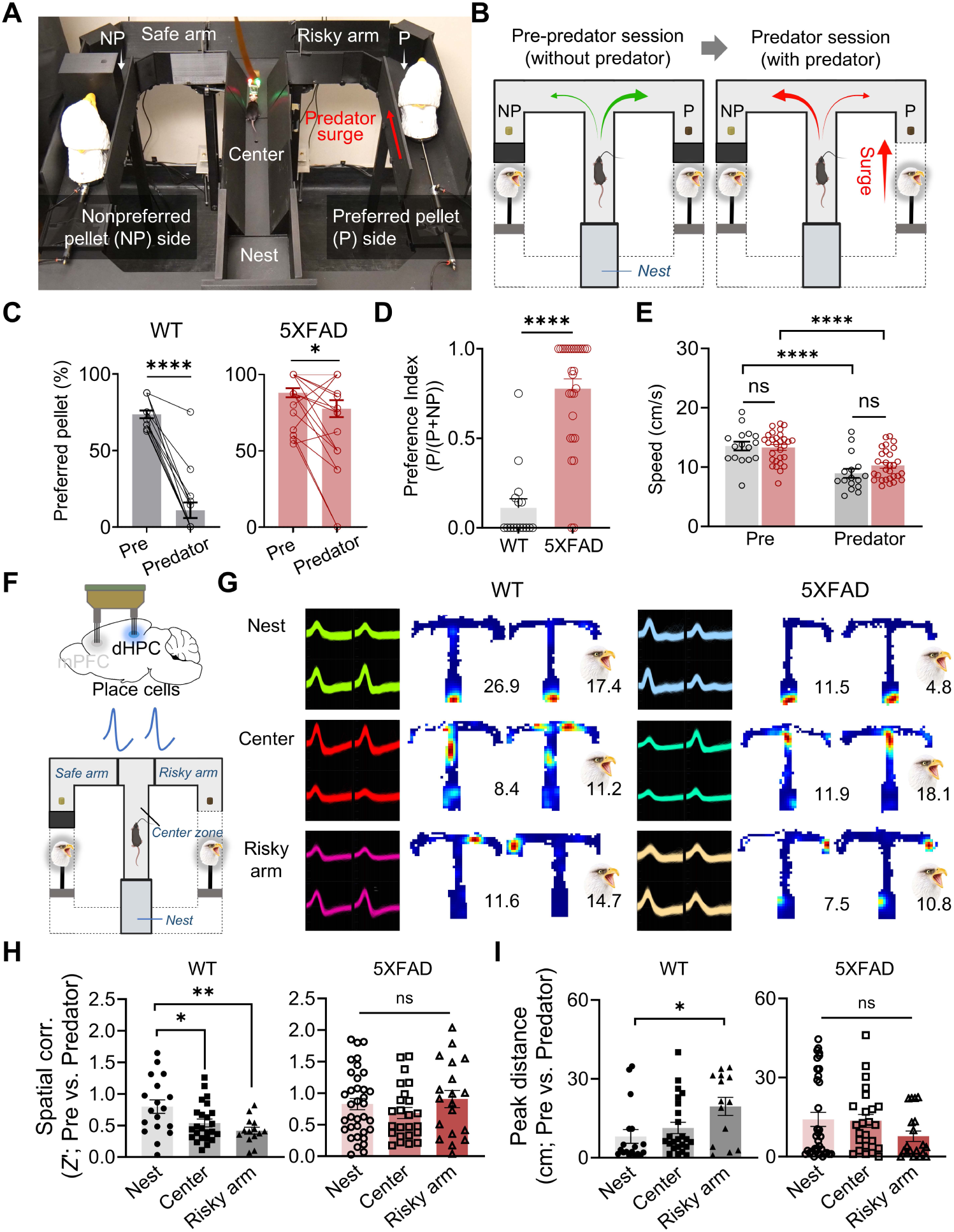
5XFAD mice exhibit inflexible foraging and rigid place cell coding in a conditional predatory task. (A) Image of a mouse tethered for neural recording in the T-maze apparatus. (B) Schematic of the behavioral procedure, consisting of successive pre-predator and predator sessions. (C) Comparison of successful procurement of preferred (P) and non-preferred (NP) pellets between pre-predator and predator sessions in WTs (6 mice, 16 recording days) and 5XFADs (11 mice, 29 recording days). (D) Preference index (ratio of preferred pellet choices to total choices) during the predator session for WT and 5XFAD mice. (E) Overall movement speed during pre-predator and predator sessions for WT and 5XFAD mice. (F) Tetrode recordings of dHPC place cells (*Top*) and designated zones in the T-maze (*Bottom*): Nest, Center zone, Risky arm, and Safe arm. (G) Representative waveforms and place fields from nest, center, and risky arm cells during pre-predator and predator sessions in WT (*Left*) and 5XFAD (*Right*) mice. The color scale (red, maximal firing; blue, no spike) corresponds to the firing rate for each unit. The numerical value indicates the peak firing rate for each session. (H,I) Spatial correlations (H) and peak distances (I) between pre-predator and predator sessions within the nest, center, and risky arm. Data are presented as mean ± SEM. \**P* < 0.05, \*\**P* < 0.01, \*\*\*\**P* < 0.0001.

When the preferred pellet arm was made “risky” by pairing it with a looming predator, WT mice adapted by shifting their foraging preference to the non-preferred, “safe” arm, thereby avoiding the predatory threat (Figure 2C). In contrast, 5XFAD mice continued to visit the predator-associated preferred pellet arm, as evidenced by significantly higher preference index (PI) values compared to WT mice (Figure 2D). Despite this divergence in foraging strategy, both groups reduced movement speed during the predator session relative to pre-predator levels, reflecting heightened caution in response to the predator. However, movement speeds did not significantly differ between groups during both pre-predator and predator sessions (Figure 2E). Histological analysis revealed higher Aβ plaque deposits in dHPC and mPFC of 5XFAD mice (Figures S2B and S2C). These findings suggest Aβ pathology impairs behavioral switching under threat while preserving baseline locomotion and food-seeking motivation.

Building on previous research showing place cell remapping in proximity to predatory threats,^44,66^ we examined how amyloid pathology affects place cell coding in 5XFAD mice (Figure 2F). Place cell activity was recorded in the dHPC of 6 WT (94 units) and 11 5XFAD (112 units) mice during pre-predator and predator sessions. Place cells were categorized by maximal firing locations during pre-predator session: nest cells (WT, n = 18; 5XFAD, n = 35), center-arm cells (WT, n = 24; 5XFAD, n = 24), risky-arm cells (WT, n = 14; 5XFAD, n = 19), and safe-arm cells (WT, n = 13; 5XFAD, n = 10) (Figures 2G, S3C, and S3D).

WT mice exhibited significant remapping of place cells in risky-arm and center-arm locations, as shown by lower spatial correlations (Z’) relative to nest cells (Figure 2H). This remapping was further supported by an increase in peak distance of place field activity in risky-arm cells compared to nest cells (Figure 2I). In contrast, 5XFAD mice showed minimal place field remapping, as indicated by spatial correlations and peak distances, suggesting impaired spatial-fear encoding and reduced neural flexibility (Figures 2H, 2I, S2D, and S2E). Place field size in risky-arm cells was larger in 5XFAD mice during the pre-predator session. However, place field size was not correlated with spatial correlation or peak distance values, indicating that the rigidity of 5XFAD place fields cannot be fully attributed to field size differences (Figures S3A-S3C).

Though a limited number of cells exceeded the visit time threshold, place field activity in the safe-arm showed spatial correlation and peak distance values comparable to those of the nest, further supporting the idea that place cells in safe areas tend to remain stable (Figures S3D and S3E). Since animals rarely visited the nonpreferred (safe) arm during the pre-predator (in most mice from both groups) and predator (in most mice from the 5XFAD group) sessions, no further analysis of place cell activity in this arm was conducted. These results show that 5XFAD mice persistently prefer dangerous zones and fail to flexibly shift behavior in response to changing threats. The corresponding lack of hippocampal place cell remapping in 5XFAD mice underscores a deficit in spatial-fear encoding, suggesting that amyloid pathology disrupts the hippocampal representation of threat-specific spatial information in dynamic, ecologically-relevant environments.

### 5XFAD mice exhibit altererd SWRs and disrupted SWR-associated dHPC activity during risky decision-making

We analyzed SWRs and dHPC neuronal activity during risky decision-making. Both 5XFAD and WT mice exhibited SWRs (Figure 3A), but 5XFAD mice showed significantly reduced dCA1 SWR frequency across all sessions (Figure 3B, Left). SWR durations were similar, with both groups showing longer SWRs during predator sessions relative to pre-predator sessions (Figure 3B, Right).

**FIGURE 3.**
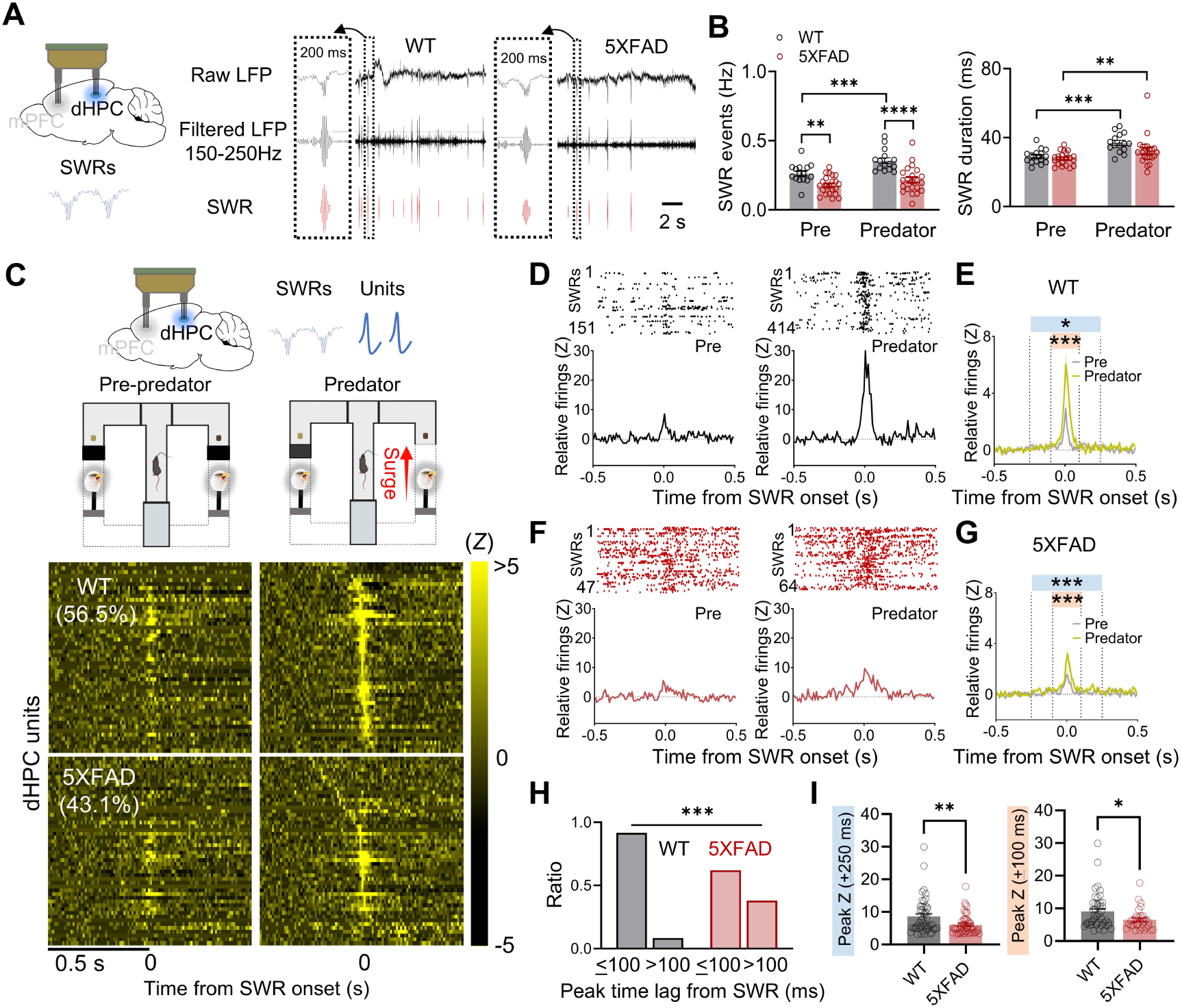
5XFAD mice exhibit alterations in SWRs and associated dHPC activity during risky decision-making. (A) Tetrode recordings of dHPC SWR activity. LFP signals were filtered (150-250 Hz) for SWR analysis. (B) Mean SWR frequency (*Left*) and duration (*Right*) in WT and 5XFAD mice across sessions. (C) *Top*: dHPC SWR and unit activity analyzed during pre-predator and predator sessions. *Bottom:* Color-coded activity (Z scores) of significantly firing dHPC units during peri-SWR epochs in predator trials, aligned to SWR onset for each session. (D) Representative WT dHPC neurons showing significant activity (z > 3) around SWR onset during the predator session. (E) Group peri-SWR activity (mean Z-scores) of WT dHPC neurons showing significant activity (z > 3) within the ±250 ms period around SWR onset during the predator session. Mean Z-scores during the ±250 ms (blue) and ±100 ms (orange) periods were compared between pre-predator and predator sessions. (F,G) Representative (F) and group (G) peri-SWR activity (mean Z-scores) of 5XFAD dHPC neurons showing significant activity (z > 3) within the ±250 ms period around SWR onset during the predator session. Mean Z-scores during the ±250 ms (blue) and ±100 ms (orange) periods were compared between pre-predator and predator sessions. (H) Proportions of dHPC neurons exhibiting significant activity within the ±250 ms vs. ±100 ms periods, compared between WT and 5XFAD groups (Chi-square test). (I) SWR-related dHPC cell activity (peak Z-scores within the ±250 ms and ±100 ms periods) during the predator session for WT and 5XFAD mice. Data are presented as mean ± SEM. \**P* < 0.05, \*\**P* < 0.01, \*\*\**P* < 0.001, \*\*\*\**P* < 0.0001.

To assess SWR modulation of dHPC firing, we analyzed peri-SWR activity (±250 ms around the SWR onset) during pre-predator and predator sessions (Figure 3C). The proportion of dHPC units exhibiting significant peri-SWR firing (z > 3) during the predator session was similar for WT (56.5%) and 5XFAD (43.1%) mice. Both groups displayed higher firing rates within the ±250 ms and ±100 ms periods during the predator session compared to pre-predator levels (Figures 3D-3G). However, key differences emerged in the temporal alignment of firing activity with SWRs. WT dHPC neurons exhibited peak firing rates closely aligned with SWR events (Figure 3H), whereas 5XFAD dHPC neurons displayed delayed peak firing relative to SWR onset (Figure S4A). Moreover, the peak firing rates of dHPC cells during the ±250 ms and ±100 ms periods were significantly higher in WT than in 5XFAD mice (Figure 3I, S4B, and S4C). These results indicate 5XFAD mice exhibit disrupted SWR-firing coordination, suggesting impaired dHPC synchronization during risky decision-making.

We next examined SWR activity and dHPC neuronal firing during a post-predator rest period within the nest (Figure S4D). Consistent with prior observations, 5XFAD mice exhibited fewer and shorter dCA1 SWRs compared to WT mice (Figure S5E). Movement speed did not differ between groups (Figure S5F), confirming SWR frequency differences were not movement-related. Analysis of peri-SWR dHPC activity (±250 ms around SWR peaks) revealed that WT mice had more SWR-modulated dHPC units and shorter SWR-firing lags than 5XFAD mice (Figures S5G and S5H). Peak firing relative to SWR onset was also delayed in 5XFAD mice (Figure S5I). However, maximal firing rates during the ±250 ms and ±100 ms periods were comparable between WT and 5XFAD (Figure S5J). These findings suggest Aβ pathology disrupts SWR-associated firing in dHPC during post-threat processing.

To examine relationships between amyloid pathology, SWR activity, and behavior, we analyzed correlations between Aβ burden, SWR frequency, and behavioral flexibility. Aβ plaque density in the dHPC and mPFC was significantly correlated with both PI values, reflecting foraging strategy (Figure S5A), and SWR frequency during task sessions (Figure S5B). Additionally, SWR frequency correlated with PI values (Figure S5C), suggesting that deficits in hippocampal SWR activity are linked to reduced behavioral flexibility in the 5XFAD mice. Our findings highlight SWR-associated hippocampal activity as a potential *in vivo* AD biomarker and its role in cognitive impairments, particularly in risky decision-making.

### 5XFAD mice exhibit altered dHPC-mPFC dynamics during risky decision-making

To examine how Aβ pathology affects dHPC-mPFC interactions, we analyzed dHPC SWRs and their influence on mPFC firing during risky decision-making (Figure 4A). Comparable proportions of mPFC neurons exhibited significant peri-SWR firing (z > 3) during predator sessions in WT (38.3%) and 5XFAD (31.9%) mice. WT mPFC cells showed increased firing within ±500 and ±250 ms of SWR onset during predator vs. pre-predator sessions (Figures 4B, S6A, and S6B), but this modulation was absent in 5XFAD mice. Mean relative mPFC firing rates in response to SWRs did not differ between pre-predator and predator sessions in 5XFAD mice (Figures 4C and S6C).

**FIGURE 4.**
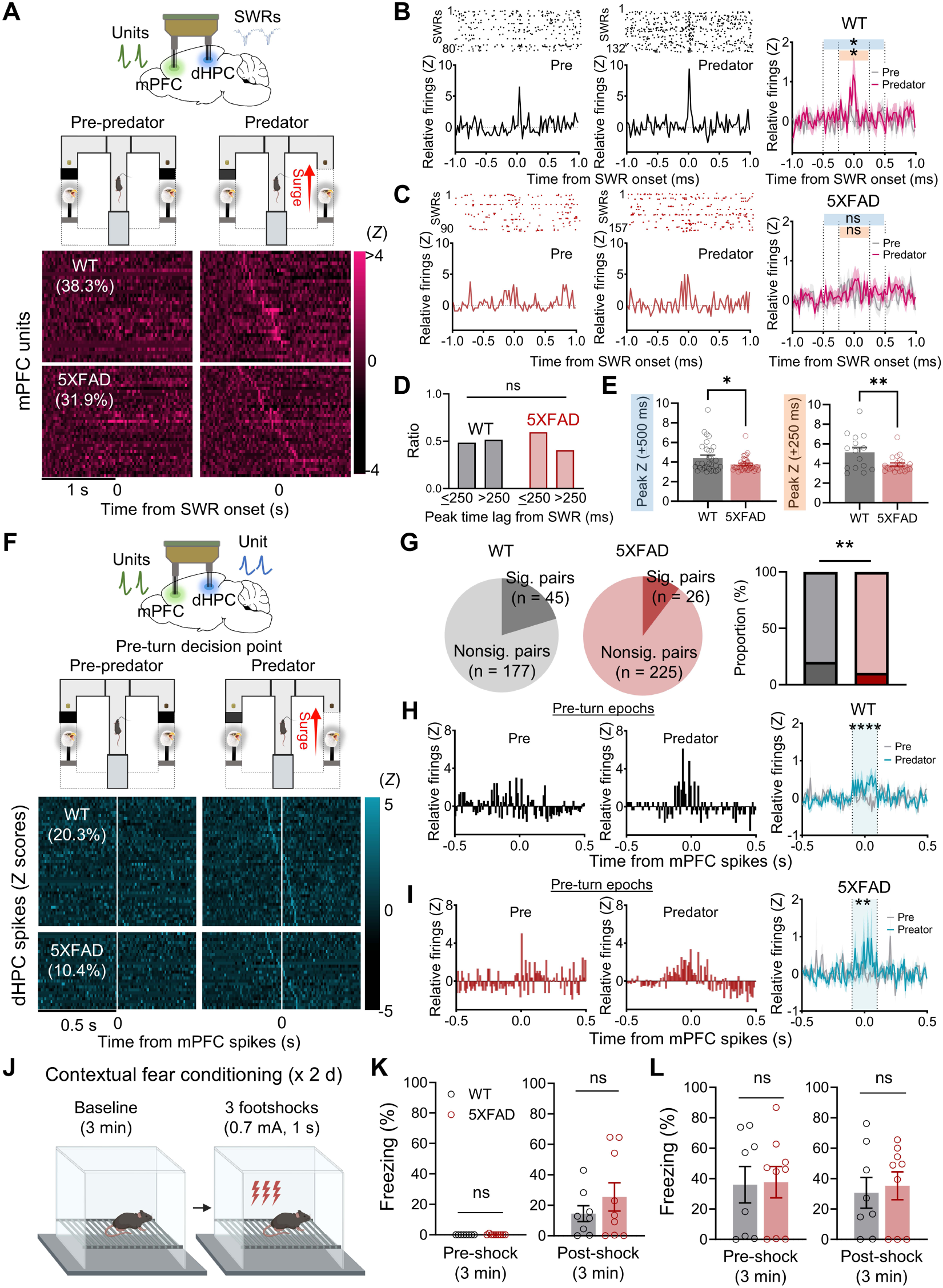
5XFAD mice exhibit altered dHPC-mPFC interactions during risky decision-making. (A) *Top:* dHPC SWR and unit activity analyzed during pre-predator and predator sessions. *Bottom:* Color-coded activity (Z scores) of significantly firing mPFC units during peri-SWR epochs in predator trials, aligned with SWR onset for each session. (B) Representative (*Left* and *Middle*) peri-SWR activity of WT mPFC neurons exhibiting significant activity (z > 3) around SWR onst during the predator session. Group peri-SWR activity (*Right*) of WT mPFC neurons exhibiting significant activity (z > 3) within the ±500 ms period around SWR onset during the predator session. Mean Z-scores during the ±500 ms (blue) and ±250 ms (orange) periods were compared between sessions. (C) Representative (*Left* and *Middle*) and group (*Right*) peri-SWR activity (mean Z-scores) of 5XFAD mPFC neurons showing significant activity (z > 3) within the ±500 ms periods around SWR onset during the predator session. Mean Z-scores during the ±500 ms and ±250 ms windows were compared between sessions. (D) Proportions of mPFC neurons exhibiting significant activity within the ±500 ms vs. ±250 ms periods in WT and 5XFAD mice. (E) Comparisons of SWR-related mPFC cell activity (peak Z-scores within the ±500 ms and ±250 ms periods) during the predator session between WT and 5XFAD mice. (F) Analysis of synchronous activity between dHPC and mPFC units during pre-predator and predator sessions. Color-coded CCs show significant dHPC-mPFC spike synchrony during 3-s pre-turn epochs before animals made turns in the presence of a predator. (G) Proportion of significant dHPC-mPFC CC pairs during the predator session. Of the available CCs, 20.3% of WT (*Left*) and 10.4% of 5XFAD (*Middle*) dHPC-mPFC pairs showed significant correlated firing. *Right:* Proportion of significant dHPC-mPFC CCs in WT and 5XFAD groups. (H,I) Representative (*Left* and *Middle*) and averaged (*Right*) dHPC-mPFC CCs from WT (H) and 5XFAD (I) mice showing significant peaks (between −100 and 100 ms; blue shaded area) during pre-turn epochs in the presence of a predator. Mean dHPC firings (Z-scores) within the ±100 ms period relative to mPFC spiking was compared between the pre-predator and predator sessions in WT and 5XFAD mice. (J) Behavioral procedure for contextual fear conditioning. (K,L) Freezing levels during pre-shock (3 min) and post-shock (3 min) phases on days 1 (K) and 2 (L) of contextual fear conditioning. Data are presented as mean ± SEM. \**P* < 0.05, \*\**P* < 0.01, \*\*\*\**P* < 0.0001.

Although the ratio of mPFC cells with peak firing near SWR onset was similar between WT and 5XFAD mice (Figure 4D), peak Z values during the ±500 ms and ±250 periods relative to SWR onset were significantly higher in WT vs. 5XFAD mPFC cells (Figure 4E). These results suggest reduced dHPC SWR-induced modulation of mPFC firing in 5XFAD mice, reflecting impaired hippocampal-prefrontal coordination during risky decision-making.

To assess decision-related dHPC-mPFC interactions, we analyzed neuronal synchrony in the ‘decision-zone’ (center arm) before animals turned toward either the preferred (risky) or non-preferred (safe) arm (Figure 4F). Cross-correlograms (CCs) were generated using mPFC cells as the reference during 3-s pre-turn epochs for pre-predator and predator decision points. We recorded 1,060 dHPC-mPFC neuronal pairs (WT: 462; 5XFAD: 598) across sessions. Of these, 222 pairs in WT and 251 in 5XFAD mice with firing rates > 0.2 Hz were analyzed.

At predator decision points, more WT dHPC-mPFC pairs (20.3%, 45/222) showed spike synchrony compared to 5XFAD pairs (10.4%, 26/251), indicating reduced dHPC-mPFC synchrony in 5XFAD mice (Figure 4G). Analysis of dHPC firing rates relative to mPFC spikes during the ±100 ms period revealed elevated mean and peak dHPC firing rates during predator vs. pre-predator sessions in both groups (Figures 4H, 4I, S6D-F). However, reduced synchrony in 5XFAD mice highlights a breakdown in dHPC-mPFC functional coordination during risky decision-making.

After the predatory task, mice underwent contextual fear conditioning over 2 days to assess their general responsiveness to aversive stimuli and learned fear behavior (Figure 4J). 5XFAD and WT mice exhibited similar levels of postshock freezing on days 1 and 2, as well as contextual freezing on day 2 (Figures 4K, 4L, S7A, and S7B). These results suggest that 5XFAD mice have normal fear learning and footshock responsiveness, indicating that altered risky foraging decisions are not due to general impairments in aversive learning or sensitivity to aversive stimuli.

Together, these findings reveal that 5XFAD mice exhibit impaired mPFC firing modulation in response to dHPC SWRs and reduced dHPC-mPFC synchrony during decision points under predatory threat. These changes suggest disruptions in hippocampal-prefrontal coordination required for adaptive decision-making, highlighting a potential mechanistic link between Aβ pathology and impaired risk-based decision-making in AD.

## DISCUSSION

This study, utilizing two distinct predatory threat scenarios and simultaneous mPFC-dHPC recordings in 5XFAD mice, reveals insights into the behavioral and neural dynamics underlying risky decision-making in AD. Behaviorally, 5XFAD mice exhibited heightened risk-taking, marked by longer foraging distances and inflexible strategies, despite normal contextual fear conditioning and threat-evoked flight responses. This selective deficit in decision-making under risk, a doman impaired in AD patients^5,8,22,67^ was observed in both 4-5 and 7-9 month-old 5XFAD mice, supporting the use of ecologically-relevant paradigms to investigate early-stage AD pathology.

At the neural level, risk-taking in 5XFAD mice coincided with reduced dCA1 SWR frequency and disrupted mPFC-dHPC connectivity. SWR frequency reductions are notable, as SWRs are established “cognitive biomarker” for memory consolidation, retrieval, and planning.^68–70^ In support, real-time disruptions and elongations of hippocampal SWRs impair and improve working memory, respectively,^54,71^ and SWRs decline with aging^72,73^ and in AD mouse models.^55,58^ Our findings linking Aβ plaques with SWR activity and risky decision-making suggests that dysfunctional SWRs play a pivotal role in 5XFAD cognitive deficits. This aligns with prior research showing the role of SWRs in spatial memory, navigation, and decision-making under risk.^52,68–71,74^

While 5XFAD and WT mice had similar SWR-associated dHPC cell firing proportions during predator encounters, 5XFAD hippocampal cells exhibited delayed peak firing relative to SWR onset. This misalignment indicates reduced synchronous firing between SWRs and hippocampal cells, consistent with prior reports of weaker hippocampal coactivation during SWRs in AD mice.^59^ Such delays likely contribute to incoherent hippocampal cell activity in 5XFAD mice.

5XFAD mice also exhibited mPFC deficits in response to dHPC SWRs, particularly under predatory threat. Predator-induced increases in mPFC firing within 500 ms of SWR onset were observed in WT but absent in 5XFAD mice. Reduced dHPC-mPFC synchrony during predator decision points further supports this notion, as fewer 5XFAD dHPC-mPFC neuronal pairs showed spike synchrony. Since mPFC neurons support rule switching, adaptive action selection, and memory-guided decisions,^53,75,76^ impaired dHPC-mPFC coordination likely underlies the inflexible foraging observed in 5XFAD mice. Strategies to restore hippocampal-prefrontal synchrony or enhance SWR activity via lifestyle interventions (e.g., exercise, diet)^77,78^ or therapeutic approaches (e.g., pharmacological,^79^ genetic,^80^ neuromodulation,^81^ or gut microbiome^82^ interventions) may mitigate cognitive decline in AD.

Place cell coding properties in Aβ mouse models show variability. For example, 16-month-old Tg2576 mice exhibit reduced spatial information and enlarged place fields,^83^ while 4-14-month-old 5XFAD mice display the opposite.^84^ Place field stability also varies across models, with 8-9-month-old 3xTg mice (which develops both Aβ and tau pathology) showing less stable place fields in circular mazes,^85^ whereas 7-13-month-old APP KI and 7-month-old J20 APP mice exhibit stable firing in familiar contexts.^86,87^ Aβ models also display experience-dependent coding deficits. APP-PS1 place cells fail to adapt to learned changes,^88^ while APP KI mice maintain stable spatial representations on a familiar linear track but struggle with remapping on a different track.^86^ Our findings of rigid place firing in 5XFAD mice, particularly in predatory threat zones, reflect impaired spatial-danger encoding, a process typically seen in normal rats^44,66^ and WT mice (Figure 2H and 2I). This deficit adds a spatial dimension to decision-making impairments in AD, highlighting an inability to integrate spatial and risk information, consistent with broader evidence of spatial memory deficits in AD.^89–91^

A couple of caveats should be considered. First, while 5XFAD mice exhibited normal contextual fear conditioning—a task engaging hippocampal function^92,93^—reduced innate fear responses may have influenced their foraging behavior. Previous studies report normal contextual fear in younger 5XFAD mice (<4 months) but impaired fear in older mice (6 months).^94^ However, we found no genotype differences in contextual freezing, even in 8-11 months-old mice. This is consistent with findings in 12-13-month-old APP-KI mice^86^ and 16-18-month-old Tg2576 mice,^95^ which also showed contextual freezing comparable to WT controls. One explanation is that contextual fear conditioning—using footshocks in a small, inescapable chamber with salient but limited cues—may mask Aβ-related hippocampal dysfunction. This aligns with Edward Thorndike’s (1899) notion that animals solve problems in constrained, unnatural environments through simple associative learning.^96^ Unlike contextual fear, foraging paradigms require complex spatial navigation and threat assessment, which may better capture AD-related cognitive deficits. Second, decision-making alterations in 5XFAD mice may stem from Aβ pathology beyond the dHPC-mPFC circuit. The amygdala, also affected in AD, integrates risk-related information into decision-making circuits.^97–99^ Given its role in threat processing and emotional learning, future studies should explore whether amygdalar neurons contribute to disruptions in dHPC-mPFC coordination and risky decision-making in AD mice.

This study highlights the value of ecologically-relevant tasks in elucidating cognitive deficits associated with AD. By simulating naturalistic decision-making, we offer a more nuanced view of how AD pathology alters behavior and neural dynamics. Notably, reliable detection of behavioral and neural alterations in 5XFAD mice with Aβ accumulations—but without tau pathology—suggests that ecological screening could detect early stage AD. This approach also enables investigations into neural mechanisms underlying early AD pathology, identifying predictive factors for AD progression, and facilitating the development of targeted interventions. Future research should explore enhancing SWR activity and hippocampal-prefrontal connectivity as strategies to prevent cognitive decline and restore cognitive function in AD.

## METHOD DETAILS

### Animals

5XFAD transgenic mice, which overexpress amyloid precursor protein (APP) and presenilin 1 (PS1), along with their WT littermate mice counterparts (aged 4-8 weeks, both sexes), were obtained from The Jackson Laboratory (stock#: 034840-JAX or 034848-JAX). Initially, these animals were group-housed in a climate-controlled vivarium under a reversed 12-h light/dark cycle (lights on at 7 PM), accredited by the Association for Assessment and Accreditation of Laboratory Animal Care (AAALAC). Upon initiation of experimental procedures, 4-9-month-old mice were individually housed and placed on a standard food-deprivation schedule, with ad libitum access to water, to maintain ∼85% of their normal body weight. A total of 17 5XFAD and 17 WT mice were assigned to the escalating predatory risk task (Figure 1), while a separate cohort (5XFAD, n=11; WT, n=6) underwent combined tetrode recording and conditional predatory risk testing (Figure 2), as detailed below. All experiments were conducted during the dark phase in strict accordance with the guidelines of the University of Washington Institutional Animal Care and Use Committee (IACUC protocol #: 4040-01).

### Surgery

Under isoflurane anesthesia, animals were secured in a stereotaxic instrument (Kopf) for the implantation of a microdrive array containing individually movable tetrodes. The tetrodes targeted the right dHPC (AP: −1.8, ML: +1.7, DV: −1.3; four tetrodes) and mPFC (AP: +1.8, ML: +0.3, DV: −1.2; four tetrodes). The tetrodes were fabricated from formvar-insulated nichrome wire (14 µm diameter; Kanthal) and gold-plated to a final impedance of 100-300 kΩ (measured at 1 kHz). The microdrive assembly was secured with dental cement and bone screws. A recovery period of at least one week preceded the initiation of behavioral experiments and recording sessions.

### Escalating predatory risk

This task was conducted in a custom-built apparatus consisting of a nest area (inner dimensions: 14 cm L × 45 cm W × 61 cm H; luminance: 2.2 lux) and an adjacent foraging space (104 cm L × 45 cm W × 61 cm H; luminance: 5.0 lux; background noise: 72-dB white noise). The environment was monitored using the ANY-maze system (Stoelting Co.) and a ceiling-mounted HD webcam (C910; Logitech) for video tracking at 30 fps (Figure 1A). The task consisted of three phases: *habituation*, *baseline foraging*, and *weasel predator testing*.

During habituation, hunger-motivated mice were acclimated by being placed in the nest with 20 food pellets (45 mg each, F0165, Bio-Serv) for 30 min/day over two days. For baseline foraging, mice were provided with two 45-mg pellets in the nest. After consuming these, they were given access to the foraging area via an open gateway to retrieve pellets placed at distances of 12.7 cm (S), 25.4 cm (M), and 38.1 cm (L) from the nest. Each daily session included three trials for four consecutive days.

The predator testing phase began with mice foraging for a pellet placed 38.1 cm from the nest in three consecutive trials. Subsequent trials introduced a taxidermy weasel mounted on a wheeled frame positioned at the opposite end of the foraging space. As the mouse approached the pellet within ∼12.7 cm, the weasel was activated to surge forward 30.4 cm towards the pellet at 60 cm/s using a pneumatic actuator, then automatically returned to its starting position. Each trial ended either when the mouse successfully secured the pellet or after 3 minutes had elapsed. This process was repeated for pellet placements at 25.2 cm and 12.7 cm.

### Conditional predatory risk

Adapted from a figure-eight maze,^100^ this T-maze setup features a nest area at the base (dimensions: 92 cm L × 92 cm W × 79 cm H; luminance: 7 lux) and two lateral goal arms, each equipped with a concealed with puppet eagle predator near a designated pellet zone (Figure 2A). Movement was tracked using headstage-mounted LEDs and integrated with the ANY-maze (Stoelting Co.) and Cheetah (NeuraLynx, Inc.) systems. This maze design ensured consistent location visits, enabling optimal place cell analysis.

The task protocol comprised four phases: *habituation*, *shaping*, *baseline foraging*, and *predator testing*, with tetrodes adjusted toward the target areas as detailed in the ‘Unit recording and analyses’ section. During habituation (two days), animals were placed in the nest for 30 min/day with ten grain (45 mg, F0165, Bio-Serv) and ten chocolate (45 mg, F0299, Bio-Serv) pellets to facilitate acclimation. The shaping phase lasted three days. Mice received one grain and one chocolate pellet in the nest. After consumption, the nest gate opened, guiding them to retrieve a pellet at increasing distances. Each arm was consistently assigned a specific pellet type. After a forced-arm retrieval and pellet consumption, the next trial began on the opposite arm. During baseline foraging, two pellets were provided in the nest. After consumption, the nest gate opened, allowing mice to choose between the left and right arm pellets. Choosing one pellet triggered the closure of the opposite arm gate. Each daily session included 6 trials, lasting for an average of 8.6 ± 1.03 days. Predator testing sessions, during which neural recordings were conducted, included *pre-predator* (8-14 trials), *predator* (3-10 trials), and *post-predator* (10 min nest period) phases. During predator trials, when mice approached the preferred pellet, a puppet eagle surged forward (15.2 cm at 38 cm/s via a linear actuator). In contrast, when mice approached the non-preferred pellet, the predator remained hidden behind a black plastic block. Pellet preference was quantified using a Preference Index (PI), defined as the ratio of preferred pellet choices to total pellet choices.

### Contextual fear conditioning

After completing the conditional predatory risk task, 5XFAD and WT mice were placed in a standard operant chamber with a stainless steel grid floor (5 mm diameter rods spaced 1 cm apart).^101^ Following a 3-minute baseline period, three unsignaled footshocks (0.7 mA, 1 sec) were administered at 1 minute intervals. Postshock freezing was assessed between each shock. The next day, mice were returned to the same chamber, and their conditioned freezing response to the context was measured for 3 minutes before administering another set of 3 unsignaled shocks and assessing postshock freezing. An uninformed observer used a custom-written computer-assisted scoring program (written in C language) to manually record the duration of freezing behavior via keystrokes on a computer keyboard.^102^ Freezing was defined as the absence of any visible movement in the body and vibrissae, except for respiratory movements.^103^ The program applied a continuous inactivity threshold of 2 seconds or more to identify freezing. The percentage of freezing was calculated by dividing the total duration of freezing by the total observation time and multiplying by 100.

### Electrophysiological recording and analysis

Single-unit and local field potential (LFP) signals were amplified (10,000X), filtered (unit: 600 Hz to 6000 Hz; LFPs: 0.1 Hz to 1000 Hz), and digitized at 32 kHz using the Cheetah data acquisition system (NeuraLynx). Unit isolation was performed using an automatic spike-sorting program (SpikeSort 3D; NeuraLynx) followed by manual cluster cutting, following protocols described in previous studies.^43,44^ Raster plots and peristimulus time histograms were generated using NeuroExplorer (Nex Technologies).

#### Place cell analysis

Place cells were identified based on criteria established in prior research.^44,66,104^ Units were classified as place cells if they exhibited stable, well-discriminated complex spike waveforms, a refractory period of at least 1 ms, peak firing rates > 2 Hz in any session, and spatial information > 0.5 bits/s in any session. Place cells were categorized according to the peak positions of their place fields during the pre-predator session as follows: *nest cells* (maximal firing in the nest), *center cells* (maximal firing in the center zone), *risky arm cells* (maximal firing in the maze arm near the threat), and *safe arm cells* (maximal firing in the maze arm away from the threat). A pixel-by-pixel spatial correlation analysis was performed using a customized R program to calculate the similarity of place maps between the pre-predator and predator sessions for each place cell. The correlation value (*r*) was transformed into a Fisher Z’ score for parametric comparisons between cell types. Additionally, the program calculated the distance between each cell’s peak firing locations across sessions. The maximal firing locations (x, y coordinates) were identified for each session, and the distance (in cm) between these points was calculated to measure positional shifts of place fields from the pre-predator to predator sessions.

#### Cross-correlogram (CC) analysis

CC of simultaneously recorded dHPC and mPFC units were generated using NeuroExplorer, following established methods.^43,66^ CCs were constructed with dHPC cells as the reference, using a 10-ms bin width for two time epochs: *pre-turn epoch_pre_* (a 3-s epoch before the animal turned at the end of the center zone toward one of the arms during the pre-predator session) and *pre-turn epoch_predator_* (a 3-s epoch before the turn during the predator session). To avoid false correlations due to covariation or nonstationary firing rates between the dHPC and mPFC, a ‘Shift-Predictor’ correction was applied, with trial shuffles performed 100 times (cf.^98^). Each shift-predictor correlogram was subtracted from its respective raw correlogram, and Z-scores were calculated using the mean and standard deviation of the corrected CCs. A neural pair was deemed significantly correlated if the peak Z-score exceeded 3. Additional criteria required that dHPC and mPFC firing rates during pre-turn periods for both the pre-predator and predator sessions be above 0.2 Hz, and the peak of the CC had to fall within a ±100 ms testing window relative to the reference spikes.

#### LFP and SWR analysis

LFP signals from a single channel of each animal’s dHPC electrodes were band-pass filtered between 150 and 250 Hz using a zero-phase digital filter to minimize phase distortions. The envelope of these high-frequency oscillations was extracted using the Hilbert transform, providing a precise measure of their amplitude. The resulting envelope was smoothed with a Gaussian-weighted moving average over a 40 ms window. SWR events were detected when the smoothed envelope exceeded a threshold of 4 standard deviations above the mean amplitude. The event ended when the amplitude fell below 2 standard deviations, ensuring precise delineation of event boundaries. Only SWR events with durations over 15 ms were included in the analysis.

#### SWR modulation of unit activity analysis

To assess the modulatory effects of SWRs on dHPC and mPFC unit activity, unit firing rates were normalized to pre-SWR baseline periods (dHPC: −500 to −250 ms; mPFC: −1000 to −500 ms). A neuron was classified as significantly activated if its peak Z-score exceeded 3 within a specified time window around SWR onset (dHPC: ±100 ms or ±250 ms; mPFC: ±250-ms or ±500 ms). All analyses were conducted using MATLAB 2024 Signal Processing Toolbox, which provided the necessary tools for detailed signal examination and processing.

#### Statistical analyses

Statistical significance was assessed using a range of tests tailored to the diverse data structures in the study. These tests included one-way ANOVAs for group comparisons, mixed-design ANOVAs for analyses incorporating both within- and between-subject variables, Pearson’s correlation, unpaired t-tests, and paired t-tests. Post hoc analyses, when required, were conducted using Tukey’s test or Fisher’s least significant difference test. The Greenhouse-Geisser correction was applied to account for sphericity violations. Comprehensive details for each statistical analysis are provided in Table S1. Statistical significance was defined as *P* < 0.05. Statistical analyses and graph generation were performed using SPSS (ver. 19), custom MATLAB codes, GraphPad Prism (ver. 9.00), and NeuroExplorer (ver. 5.030).

#### Histology

After concluding the experiment, electrolytic lesions were created by applying a 10 µA current for 10 s to the tetrode tips to mark electrode placements. Mice were then euthanized with an overdose of Beuthanasia and perfused intracardially with phosphate-buffered saline (PBS), followed by fixation with 4% paraformaldehyde in PBS. Extracted brains were stored overnight at 4 °C in fixative, then transferred to 10%, 20%, and 30% sucrose solutions until they sank. For Aβ immunostaining, transverse brain sections (30 µm) were rinsed in PBS and incubated for 1 hr in 5% normal goat serum with 0.5% TritonX in PBS to block non-specific binding. The sections were then incubated overnight at 4 °C with Aβ-specific mouse monoclonal antibody (1:1000, MediMabs, McSA1). After thorough washing, sections were incubated with a goat anti-mouse secondary antibody conjugated to Alexa Fluor 568 (1:250, Abcam, ab175473) for 2 hr at room temperature. The prepared sections were mounted on slides and cover slipped with Flouromount-G^TM^ containing DAPI (eBioscience) for nuclear staining.

Imaging was performed using a Keyence BZ-X800E microscope. Image analysis was conducted using ImageJ software (NIH, version 1.54d). A separate set of brain sections was mounted onto gelatinized slides and stained with Cresyl violet and Prussian blue to confirm the precise locations of the tetrode tips.

## ACKNOWLEGMENTS

This study was supported by the National Institutes of Health grants AG067008 (E.J.K.), MH099073 (J.J.K.), the University of Washington Royalty Research Fund A188443 (E.J.K.), and the Ministry of Science and ICT through the National Research Foundation of Korea: Brain Science Research Program NRF-2022M3E5E8018421 and NRF-2022R1A2C2009265 (J.C.). We thank Drs. Min Whan Jung and Jong Won Lee from the Korea Advanced Institute of Science and Technology for generously sharing custom-made microdrive parts and electronic interface boards, and Nayoung Kim for assistance with experiments and data collection/analysis.

## AUTHOR CONTRIBUTIONS

E.J.K., J.C., and J.J.K. designed the research; E.J.K., S.G.P., B.P.S., and H.B. performed behavioral and neurophysiological experiments and analyses; and E.J.K., S.G.P., J.C., and J.J.K. wrote the manuscript.

## SUPPLEMENTARY INFORMATION AND DATA AVAILABILITY

The datasets and analysis code for this research are available in the Dryad data repository upon manuscript acceptance.

## REFERENCES

1. Bangma, D.F., Tucha, O., Tucha, L., De Deyn, P.P., and Koerts, J. (2021). How well do people living with neurodegenerative diseases manage their finances? A meta-analysis and systematic review on the capacity to make financial decisions in people living with neurodegenerative diseases. Neurosci Biobehav Rev 127, 709–739. 10.1016/j.neubiorev.2021.05.021.

2. Bertrand, E., van Duinkerken, E., Landeira-Fernandez, J., Dourado, M.C.N., Santos, R.L., Laks, J., and Mograbi, D.C. (2017). Behavioral and Psychological Symptoms Impact Clinical Competence in Alzheimer’s Disease. Front Aging Neurosci 9, 182. 10.3389/fnagi.2017.00182.

3. Delazer, M., Sinz, H., Zamarian, L., and Benke, T. (2007). Decision-making with explicit and stable rules in mild Alzheimer’s disease. Neuropsychologia 45, 1632–1641. 10.1016/j.neuropsychologia.2007.01.006.

4. Gleichgerrcht, E., Ibanez, A., Roca, M., Torralva, T., and Manes, F. (2010). Decision-making cognition in neurodegenerative diseases. Nat Rev Neurol 6, 611–623. 10.1038/nrneurol.2010.148.

5. Ha, J., Kim, E.J., Lim, S., Shin, D.W., Kang, Y.J., Bae, S.M., Yoon, H.K., and Oh, K.S. (2012). Altered risk-aversion and risk-taking behaviour in patients with Alzheimer’s disease. Psychogeriatrics 12, 151–158. 10.1111/j.1479-8301.2011.00396.x.

6. Marson, D.C., Martin, R.C., Wadley, V., Griffith, H.R., Snyder, S., Goode, P.S., Kinney, F.C., Nicholas, A.P., Steele, T., Anderson, B., et al. (2009). Clinical interview assessment of financial capacity in older adults with mild cognitive impairment and Alzheimer’s disease. J Am Geriatr Soc 57, 806–814. 10.1111/j.1532-5415.2009.02202.x.

7. Okonkwo, O., Griffith, H.R., Belue, K., Lanza, S., Zamrini, E.Y., Harrell, L.E., Brockington, J.C., Clark, D., Raman, R., and Marson, D.C. (2007). Medical decision-making capacity in patients with mild cognitive impairment. Neurology 69, 1528–1535. 10.1212/01.wnl.0000277639.90611.d9.

8. Sun, T., Xie, T., Wang, J., Zhang, L., Tian, Y., Wang, K., Yu, X., and Wang, H. (2020). Decision-Making Under Ambiguity or Risk in Individuals With Alzheimer’s Disease and Mild Cognitive Impairment. Front Psychiatry 11, 218. 10.3389/fpsyt.2020.00218.

9. Sun, W., Matsuoka, T., and Narumoto, J. (2021). Decision-Making Support for People With Alzheimer’s Disease: A Narrative Review. Front Psychol 12, 750803. 10.3389/fpsyg.2021.750803.

10. van Duinkerken, E., Farme, J., Landeira-Fernandez, J., Dourado, M.C., Laks, J., and Mograbi, D.C. (2018). Medical and Research Consent Decision-Making Capacity in Patients with Alzheimer’s Disease: A Systematic Review. J Alzheimers Dis 65, 917–930. 10.3233/JAD-180311.

11. Boyle, P.A., Yu, L., Schneider, J.A., Wilson, R.S., and Bennett, D.A. (2019). Scam Awareness Related to Incident Alzheimer Dementia and Mild Cognitive Impairment: A Prospective Cohort Study. Ann Intern Med 170, 702–709. 10.7326/M18-2711.

12. Davis, R., Ziomkowski, M.K., and Veltkamp, A. (2017). Everyday Decision Making in Individuals with Early-Stage Alzheimer’s Disease: An Integrative Review of the Literature. Res Gerontol Nurs 10, 240–247. 10.3928/19404921-20170831-05.

13. Gerstenecker, A., Triebel, K.L., Martin, R., Snyder, S., and Marson, D.C. (2016). Both Financial and Cognitive Decline Predict Clinical Progression in MCI. Alzheimer Dis Assoc Disord 30, 27–34. 10.1097/WAD.0000000000000120.

14. Han, S.D., Boyle, P.A., James, B.D., Yu, L., and Bennett, D.A. (2015). Mild cognitive impairment is associated with poorer decision-making in community-based older persons. J Am Geriatr Soc 63, 676–683. 10.1111/jgs.13346.

15. Martin, R.C., Gerstenecker, A., Triebel, K.L., Falola, M., McPherson, T., Cutter, G., and Marson, D.C. (2019). Declining Financial Capacity in Mild Cognitive Impairment: A Six-Year Longitudinal Study. Arch Clin Neuropsychol 34, 152–161. 10.1093/arclin/acy030.

16. Niccolai, L.M., Triebel, K.L., Gerstenecker, A., McPherson, T.O., Cutter, G.R., Martin, R.C., and Marson, D.C. (2017). Neurocognitive Predictors of Declining Financial Capacity in Persons with Mild Cognitive Impairment. Clin Gerontol 40, 14–23. 10.1080/07317115.2016.1228022.

17. Okonkwo, O.C., Griffith, H.R., Copeland, J.N., Belue, K., Lanza, S., Zamrini, E.Y., Harrell, L.E., Brockington, J.C., Clark, D., Raman, R., and Marson, D.C. (2008). Medical decision-making capacity in mild cognitive impairment: a 3-year longitudinal study. Neurology 71, 1474–1480. 10.1212/01.wnl.0000334301.32358.48.

18. Spreng, R.N.P., Karlawish, J.M., and Marson, D.C.M. (2016). Cognitive, social, and neural determinants of diminished decision-making and financial exploitation risk in aging and dementia: A review and new model. J Elder Abuse Negl 28, 320–344. 10.1080/08946566.2016.1237918.

19. Triebel, K.L., Martin, R., Griffith, H.R., Marceaux, J., Okonkwo, O.C., Harrell, L., Clark, D., Brockington, J., Bartolucci, A., and Marson, D.C. (2009). Declining financial capacity in mild cognitive impairment: A 1-year longitudinal study. Neurology 73, 928–934. 10.1212/WNL.0b013e3181b87971.

20. Yerstein, O., Carr, A.R., Jimenez, E., and Mendez, M.F. (2020). Neuropsychiatric Effects on Decision-Making in Early Alzheimer Disease. J Geriatr Psychiatry Neurol 33, 68–72. 10.1177/0891988719888292.

21. Kapasi, A., Yu, L., Stewart, C., Schneider, J.A., Bennett, D.A., and Boyle, P.A. (2021). Association of Amyloid-beta Pathology with Decision Making and Scam Susceptibility. J Alzheimers Dis 83, 879–887. 10.3233/JAD-210356.

22. Sinz, H., Zamarian, L., Benke, T., Wenning, G.K., and Delazer, M. (2008). Impact of ambiguity and risk on decision making in mild Alzheimer’s disease. Neuropsychologia 46, 2043–2055. 10.1016/j.neuropsychologia.2008.02.002.

23. Grossberg, S., and Gutowski, W.E. (1987). Neural dynamics of decision making under risk: affective balance and cognitive-emotional interactions. Psychol Rev 94, 300–318.

24. Heilman, R.M., Crisan, L.G., Houser, D., Miclea, M., and Miu, A.C. (2010). Emotion regulation and decision making under risk and uncertainty. Emotion 10, 257–265. 10.1037/a0018489.

25. Wang, Y., and Ruhe, G. (2007). The cognitive process of decision making. International Journal of Cognitive Informatics and Natural Intelligence 1, 73–85.

26. Anderson, R., Hart, D.W., Sweis, B., Sherman, M.A., Thomas, M.J., Redish, A.D., and Lesne, S.E. (2021). Sex-specific interactions between procedural and deliberative decision-making systems in a mouse model of Alzheimer’s disease. bioRxiv. 10.1101/2021.03.16.435712.

27. Bateman, R.J., Smith, J., Donohue, M.C., Delmar, P., Abbas, R., Salloway, S., Wojtowicz, J., Blennow, K., Bittner, T., Black, S.E., et al. (2023). Two Phase 3 Trials of Gantenerumab in Early Alzheimer’s Disease. N Engl J Med 389, 1862–1876. 10.1056/NEJMoa2304430.

28. Cacabelos, R. (2018). Have there been improvements in Alzheimer’s disease drug discovery over the past 5 years? Expert Opin Drug Discov 13, 523–538. 10.1080/17460441.2018.1457645.

29. Mehta, D., Jackson, R., Paul, G., Shi, J., and Sabbagh, M. (2017). Why do trials for Alzheimer’s disease drugs keep failing? A discontinued drug perspective for 2010-2015. Expert Opin Investig Drugs 26, 735–739. 10.1080/13543784.2017.1323868.

30. Driscoll, I., and Troncoso, J. (2011). Asymptomatic Alzheimer’s disease: a prodrome or a state of resilience? Curr Alzheimer Res 8, 330–335. 10.2174/156720511795745348.

31. Hohman, T.J., McLaren, D.G., Mormino, E.C., Gifford, K.A., Libon, D.J., Jefferson, A.L., and Alzheimer’s Disease Neuroimaging, I. (2016). Asymptomatic Alzheimer disease: Defining resilience. Neurology 87, 2443–2450. 10.1212/WNL.0000000000003397.

32. Latimer, C.S., Keene, C.D., Flanagan, M.E., Hemmy, L.S., Lim, K.O., White, L.R., Montine, K.S., and Montine, T.J. (2017). Resistance to Alzheimer Disease Neuropathologic Changes and Apparent Cognitive Resilience in the Nun and Honolulu-Asia Aging Studies. J Neuropathol Exp Neurol 76, 458–466. 10.1093/jnen/nlx030.

33. Boyle, R., Klinger, H.M., Shirzadi, Z., Coughlan, G.T., Seto, M., Properzi, M.J., Townsend, D.L., Yuan, Z., Scanlon, C., Jutten, R.J., et al. (2024). Left Frontoparietal Control Network Connectivity Moderates the Effect of Amyloid on Cognitive Decline in Preclinical Alzheimer’s Disease: The A4 Study. J Prev Alzheimers Dis 11, 881–888. 10.14283/jpad.2024.140.

34. Buckley, R.F., Schultz, A.P., Hedden, T., Papp, K.V., Hanseeuw, B.J., Marshall, G., Sepulcre, J., Smith, E.E., Rentz, D.M., Johnson, K.A., et al. (2017). Functional network integrity presages cognitive decline in preclinical Alzheimer disease. Neurology 89, 29–37. 10.1212/WNL.0000000000004059.

35. Walsh, C., Drinkenburg, W.H., and Ahnaou, A. (2017). Neurophysiological assessment of neural network plasticity and connectivity: Progress towards early functional biomarkers for disease interception therapies in Alzheimer’s disease. Neurosci Biobehav Rev 73, 340–358. 10.1016/j.neubiorev.2016.12.020.

36. Corcoran, K.A., and Maren, S. (2001). Hippocampal inactivation disrupts contextual retrieval of fear memory after extinction. J Neurosci 21, 1720–1726.

37. Corcoran, K.A., and Quirk, G.J. (2007). Activity in prelimbic cortex is necessary for the expression of learned, but not innate, fears. J Neurosci 27, 840–844. 10.1523/JNEUROSCI.5327-06.2007.

38. Gluth, S., Sommer, T., Rieskamp, J., and Buchel, C. (2015). Effective Connectivity between Hippocampus and Ventromedial Prefrontal Cortex Controls Preferential Choices from Memory. Neuron 86, 1078–1090. 10.1016/j.neuron.2015.04.023.

39. Rhodes, S.E., and Murray, E.A. (2013). Differential effects of amygdala, orbital prefrontal cortex, and prelimbic cortex lesions on goal-directed behavior in rhesus macaques. J Neurosci 33, 3380–3389. 10.1523/JNEUROSCI.4374-12.2013.

40. Yoon, T., Okada, J., Jung, M.W., and Kim, J.J. (2008). Prefrontal cortex and hippocampus subserve different components of working memory in rats. Learn Mem 15, 97–105. 10.1101/lm.850808.

41. Yu, J.Y., and Frank, L.M. (2015). Hippocampal-cortical interaction in decision making. Neurobiol Learn Mem 117, 34–41. 10.1016/j.nlm.2014.02.002.

42. Euston, D.R., Gruber, A.J., and McNaughton, B.L. (2012). The role of medial prefrontal cortex in memory and decision making. Neuron 76, 1057–1070. 10.1016/j.neuron.2012.12.002.

43. Kim, E.J., Kong, M.S., Park, S.G., Mizumori, S.J.Y., Cho, J., and Kim, J.J. (2018). Dynamic coding of predatory information between the prelimbic cortex and lateral amygdala in foraging rats. Sci Adv 4, eaar7328. 10.1126/sciadv.aar7328.

44. Kim, E.J., Park, M., Kong, M.S., Park, S.G., Cho, J., and Kim, J.J. (2015). Alterations of hippocampal place cells in foraging rats facing a “predatory” threat. Curr Biol 25, 1362–1367. 10.1016/j.cub.2015.03.048.

45. Bayard, S., Jacus, J.P., Raffard, S., and Gely-Nargeot, M.C. (2014). Apathy and emotion-based decision-making in amnesic mild cognitive impairment and Alzheimer’s disease. Behav Neurol 2014, 231469. 10.1155/2014/231469.

46. Eimer, W.A., and Vassar, R. (2013). Neuron loss in the 5XFAD mouse model of Alzheimer’s disease correlates with intraneuronal Abeta42 accumulation and Caspase-3 activation. Mol Neurodegener 8, 2. 10.1186/1750-1326-8-2.

47. Oakley, H., Cole, S.L., Logan, S., Maus, E., Shao, P., Craft, J., Guillozet-Bongaarts, A., Ohno, M., Disterhoft, J., Van Eldik, L., et al. (2006). Intraneuronal beta-amyloid aggregates, neurodegeneration, and neuron loss in transgenic mice with five familial Alzheimer’s disease mutations: potential factors in amyloid plaque formation. J Neurosci 26, 10129–10140. 10.1523/JNEUROSCI.1202-06.2006.

48. Girard, S.D., Baranger, K., Gauthier, C., Jacquet, M., Bernard, A., Escoffier, G., Marchetti, E., Khrestchatisky, M., Rivera, S., and Roman, F.S. (2013). Evidence for early cognitive impairment related to frontal cortex in the 5XFAD mouse model of Alzheimer’s disease. J Alzheimers Dis 33, 781–796. 10.3233/JAD-2012-120982.

49. Sadleir, K.R., Popovic, J., and Vassar, R. (2018). ER stress is not elevated in the 5XFAD mouse model of Alzheimer’s disease. J Biol Chem 293, 18434–18443. 10.1074/jbc.RA118.005769.

50. Girardeau, G., Inema, I., and Buzsaki, G. (2017). Reactivations of emotional memory in the hippocampus-amygdala system during sleep. Nat Neurosci 20, 1634–1642. 10.1038/nn.4637.

51. Schmidt, B., and Redish, A.D. (2021). Disrupting the medial prefrontal cortex with designer receptors exclusively activated by designer drug alters hippocampal sharp-wave ripples and their associated cognitive processes. Hippocampus 31, 1051–1067. 10.1002/hipo.23367.

52. Colgin, L.L. (2011). Oscillations and hippocampal-prefrontal synchrony. Curr Opin Neurobiol 21, 467–474. 10.1016/j.conb.2011.04.006.

53. den Bakker, H., Van Dijck, M., Sun, J.J., and Kloosterman, F. (2023). Sharp-wave-ripple-associated activity in the medial prefrontal cortex supports spatial rule switching. Cell Rep 42, 112959. 10.1016/j.celrep.2023.112959.

54. Jadhav, S.P., Rothschild, G., Roumis, D.K., and Frank, L.M. (2016). Coordinated Excitation and Inhibition of Prefrontal Ensembles during Awake Hippocampal Sharp-Wave Ripple Events. Neuron 90, 113–127. 10.1016/j.neuron.2016.02.010.

55. Jura, B., Macrez, N., Meyrand, P., and Bem, T. (2019). Deficit in hippocampal ripples does not preclude spatial memory formation in APP/PS1 mice. Sci Rep 9, 20129. 10.1038/s41598-019-56582-w.

56. Gillespie, A.K., Jones, E.A., Lin, Y.H., Karlsson, M.P., Kay, K., Yoon, S.Y., Tong, L.M., Nova, P., Carr, J.S., Frank, L.M., and Huang, Y. (2016). Apolipoprotein E4 Causes Age-Dependent Disruption of Slow Gamma Oscillations during Hippocampal Sharp-Wave Ripples. Neuron 90, 740–751. 10.1016/j.neuron.2016.04.009.

57. Jones, E.A., Gillespie, A.K., Yoon, S.Y., Frank, L.M., and Huang, Y. (2019). Early Hippocampal Sharp-Wave Ripple Deficits Predict Later Learning and Memory Impairments in an Alzheimer’s Disease Mouse Model. Cell Rep 29, 2123–2133 e2124. 10.1016/j.celrep.2019.10.056.

58. Iaccarino, H.F., Singer, A.C., Martorell, A.J., Rudenko, A., Gao, F., Gillingham, T.Z., Mathys, H., Seo, J., Kritskiy, O., Abdurrob, F., et al. (2016). Gamma frequency entrainment attenuates amyloid load and modifies microglia. Nature 540, 230–235. 10.1038/nature20587.

59. Prince, S.M., Paulson, A.L., Jeong, N., Zhang, L., Amigues, S., and Singer, A.C. (2021). Alzheimer’s pathology causes impaired inhibitory connections and reactivation of spatial codes during spatial navigation. Cell Rep 35, 109008. 10.1016/j.celrep.2021.109008.

60. Krakauer, J.W., Ghazanfar, A.A., Gomez-Marin, A., MacIver, M.A., and Poeppel, D. (2017). Neuroscience Needs Behavior: Correcting a Reductionist Bias. Neuron 93, 480–490. 10.1016/j.neuron.2016.12.041.

61. Kempermann, G., Kuhn, H.G., and Gage, F.H. (1997). More hippocampal neurons in adult mice living in an enriched environment. Nature 386, 493–495. 10.1038/386493a0.

62. Bennett, E.L., Diamond, M.C., Krech, D., and Rosenzweig, M.R. (1964). Chemical and Anatomical Plasticity Brain. Science 146, 610–619. 10.1126/science.146.3644.610.

63. Pellman, B.A., and Kim, J.J. (2016). What Can Ethobehavioral Studies Tell Us about the Brain’s Fear System? Trends Neurosci 39, 420–431. 10.1016/j.tins.2016.04.001.

64. Choi, J.S., and Kim, J.J. (2010). Amygdala regulates risk of predation in rats foraging in a dynamic fear environment. Proc Natl Acad Sci U S A 107, 21773–21777. 10.1073/pnas.1010079108.

65. Kim, J.J., Choi, J.-S., and Lee, H.J. (2016). Foraging in the face of fear: novel strategies for evaluating amygdala functions in rats. In Living Without an Amygdala, D.G. Amaral, and R. Adolphs, eds. (The Guilford Press).

66. Kong, M.S., Kim, E.J., Park, S., Zweifel, L.S., Huh, Y., Cho, J., and Kim, J.J. (2021). ’Fearful-place’ coding in the amygdala-hippocampal network. Elife 10. 10.7554/eLife.72040.

67. Gorrie, C.A., Brown, J., and Waite, P.M. (2008). Crash characteristics of older pedestrian fatalities: dementia pathology may be related to ‘at risk’ traffic situations. Accid Anal Prev 40, 912–919. 10.1016/j.aap.2007.10.006.

68. Buzsaki, G. (2015). Hippocampal sharp wave-ripple: A cognitive biomarker for episodic memory and planning. Hippocampus 25, 1073–1188. 10.1002/hipo.22488.

69. Joo, H.R., and Frank, L.M. (2018). The hippocampal sharp wave-ripple in memory retrieval for immediate use and consolidation. Nat Rev Neurosci 19, 744–757. 10.1038/s41583-018-0077-1.

70. Norman, Y., Yeagle, E.M., Khuvis, S., Harel, M., Mehta, A.D., and Malach, R. (2019). Hippocampal sharp-wave ripples linked to visual episodic recollection in humans. Science 365. 10.1126/science.aax1030.

71. Fernandez-Ruiz, A., Oliva, A., Fermino de Oliveira, E., Rocha-Almeida, F., Tingley, D., and Buzsaki, G. (2019). Long-duration hippocampal sharp wave ripples improve memory. Science 364, 1082–1086. 10.1126/science.aax0758.

72. Cowen, S.L., Gray, D.T., Wiegand, J.L., Schimanski, L.A., and Barnes, C.A. (2020). Age-associated changes in waking hippocampal sharp-wave ripples. Hippocampus 30, 28–38. 10.1002/hipo.23005.

73. Wiegand, J.P., Gray, D.T., Schimanski, L.A., Lipa, P., Barnes, C.A., and Cowen, S.L. (2016). Age Is Associated with Reduced Sharp-Wave Ripple Frequency and Altered Patterns of Neuronal Variability. J Neurosci 36, 5650–5660. 10.1523/JNEUROSCI.3069-15.2016.

74. Papale, A.E., Zielinski, M.C., Frank, L.M., Jadhav, S.P., and Redish, A.D. (2016). Interplay between Hippocampal Sharp-Wave-Ripple Events and Vicarious Trial and Error Behaviors in Decision Making. Neuron 92, 975–982. 10.1016/j.neuron.2016.10.028.

75. Jarovi, J., Pilkiw, M., and Takehara-Nishiuchi, K. (2023). Prefrontal neuronal ensembles link prior knowledge with novel actions during flexible action selection. Cell Rep 42, 113492. 10.1016/j.celrep.2023.113492.

76. Tang, W., Shin, J.D., Frank, L.M., and Jadhav, S.P. (2017). Hippocampal-Prefrontal Reactivation during Learning Is Stronger in Awake Compared with Sleep States. J Neurosci 37, 11789–11805. 10.1523/JNEUROSCI.2291-17.2017.

77. Morris, J.K., Vidoni, E.D., Johnson, D.K., Van Sciver, A., Mahnken, J.D., Honea, R.A., Wilkins, H.M., Brooks, W.M., Billinger, S.A., Swerdlow, R.H., and Burns, J.M. (2017). Aerobic exercise for Alzheimer’s disease: A randomized controlled pilot trial. PLoS One 12, e0170547. 10.1371/journal.pone.0170547.

78. Scarmeas, N., Stern, Y., Tang, M.X., Mayeux, R., and Luchsinger, J.A. (2006). Mediterranean diet and risk for Alzheimer’s disease. Ann Neurol 59, 912–921. 10.1002/ana.20854.

79. Mangialasche, F., Solomon, A., Winblad, B., Mecocci, P., and Kivipelto, M. (2010). Alzheimer’s disease: clinical trials and drug development. Lancet Neurol 9, 702–716. 10.1016/S1474-4422(10)70119-8.

80. Tuszynski, M.H., Thal, L., Pay, M., Salmon, D.P., U, H.S., Bakay, R., Patel, P., Blesch, A., Vahlsing, H.L., Ho, G., et al. (2005). A phase 1 clinical trial of nerve growth factor gene therapy for Alzheimer disease. Nat Med 11, 551–555. 10.1038/nm1239.

81. Pople, C.B., Meng, Y., Li, D.Z., Bigioni, L., Davidson, B., Vecchio, L.M., Hamani, C., Rabin, J.S., and Lipsman, N. (2020). Neuromodulation in the Treatment of Alzheimer’s Disease: Current and Emerging Approaches. J Alzheimers Dis 78, 1299–1313. 10.3233/JAD-200913.

82. Vogt, N.M., Kerby, R.L., Dill-McFarland, K.A., Harding, S.J., Merluzzi, A.P., Johnson, S.C., Carlsson, C.M., Asthana, S., Zetterberg, H., Blennow, K., et al. (2017). Gut microbiome alterations in Alzheimer’s disease. Sci Rep 7, 13537. 10.1038/s41598-017-13601-y.

83. Cacucci, F., Yi, M., Wills, T.J., Chapman, P., and O’Keefe, J. (2008). Place cell firing correlates with memory deficits and amyloid plaque burden in Tg2576 Alzheimer mouse model. Proc Natl Acad Sci U S A 105, 7863–7868. 10.1073/pnas.0802908105.

84. Zhang, H., Chen, L., Johnston, K.G., Crapser, J., Green, K.N., Ha, N.M., Tenner, A.J., Holmes, T.C., Nitz, D.A., and Xu, X. (2023). Degenerate mapping of environmental location presages deficits in object-location encoding and memory in the 5xFAD mouse model for Alzheimer’s disease. Neurobiol Dis 176, 105939. 10.1016/j.nbd.2022.105939.

85. Mably, A.J., Gereke, B.J., Jones, D.T., and Colgin, L.L. (2017). Impairments in spatial representations and rhythmic coordination of place cells in the 3xTg mouse model of Alzheimer’s disease. Hippocampus 27, 378–392. 10.1002/hipo.22697.

86. Jun, H., Bramian, A., Soma, S., Saito, T., Saido, T.C., and Igarashi, K.M. (2020). Disrupted Place Cell Remapping and Impaired Grid Cells in a Knockin Model of Alzheimer’s Disease. Neuron 107, 1095–1112 e1096. 10.1016/j.neuron.2020.06.023.

87. Ying, J., Keinath, A.T., Lavoie, R., Vigneault, E., El Mestikawy, S., and Brandon, M.P. (2022). Disruption of the grid cell network in a mouse model of early Alzheimer’s disease. Nat Commun 13, 886. 10.1038/s41467-022-28551-x.

88. Cayzac, S., Mons, N., Ginguay, A., Allinquant, B., Jeantet, Y., and Cho, Y.H. (2015). Altered hippocampal information coding and network synchrony in APP-PS1 mice. Neurobiol Aging 36, 3200–3213. 10.1016/j.neurobiolaging.2015.08.023.

89. Elder, G.A., Gama Sosa, M.A., and De Gasperi, R. (2010). Transgenic mouse models of Alzheimer’s disease. Mt Sinai J Med 77, 69–81. 10.1002/msj.20159.

90. Gotz, J., and Ittner, L.M. (2008). Animal models of Alzheimer’s disease and frontotemporal dementia. Nat Rev Neurosci 9, 532–544. 10.1038/nrn2420.

91. Jankowsky, J.L., and Zheng, H. (2017). Practical considerations for choosing a mouse model of Alzheimer’s disease. Mol Neurodegener 12, 89. 10.1186/s13024-017-0231-7.

92. Kim, J.J., and Fanselow, M.S. (1992). Modality-specific retrograde amnesia of fear. Science 256, 675–677. 10.1126/science.1585183.

93. Phillips, R.G., and LeDoux, J.E. (1992). Differential contribution of amygdala and hippocampus to cued and contextual fear conditioning. Behav Neurosci 106, 274–285. 10.1037//0735-7044.106.2.274.

94. Kimura, R., and Ohno, M. (2009). Impairments in remote memory stabilization precede hippocampal synaptic and cognitive failures in 5XFAD Alzheimer mouse model. Neurobiol Dis 33, 229–235. 10.1016/j.nbd.2008.10.006.

95. Corcoran, K.A., Lu, Y., Turner, R.S., and Maren, S. (2002). Overexpression of hAPPswe impairs rewarded alternation and contextual fear conditioning in a transgenic mouse model of Alzheimer’s disease. Learn Mem 9, 243–252. 10.1101/lm.51002.

96. Thorndike, E. (1899). Instinct. In Biological lectures from the Marine Laboratory at Woods Holl, (Ginn & Company), pp. 57–67.

97. Bechara, A., Damasio, H., and Damasio, A.R. (2003). Role of the amygdala in decision-making. Ann N Y Acad Sci 985, 356–369. 10.1111/j.1749-6632.2003.tb07094.x.

98. Burgos-Robles, A., Kimchi, E.Y., Izadmehr, E.M., Porzenheim, M.J., Ramos-Guasp, W.A., Nieh, E.H., Felix-Ortiz, A.C., Namburi, P., Leppla, C.A., Presbrey, K.N., et al. (2017). Amygdala inputs to prefrontal cortex guide behavior amid conflicting cues of reward and punishment. Nat Neurosci 20, 824–835. 10.1038/nn.4553.

99. Seymour, B., and Dolan, R. (2008). Emotion, decision making, and the amygdala. Neuron 58, 662–671. 10.1016/j.neuron.2008.05.020.

100. Pedigo, S.F., Song, E.Y., Jung, M.W., and Kim, J.J. (2006). A computer vision-based automated Figure-8 maze for working memory test in rodents. J Neurosci Methods 156, 10–16. 10.1016/j.jneumeth.2006.01.029.

101. Chen, C., Kim, J.J., Thompson, R.F., and Tonegawa, S. (1996). Hippocampal lesions impair contextual fear conditioning in two strains of mice. Behav Neurosci 110, 1177–1180.

102. Kim, E.J., Kim, E.S., Covey, E., and Kim, J.J. (2010). Social transmission of fear in rats: the role of 22-kHz ultrasonic distress vocalization. PLoS One 5, e15077. 10.1371/journal.pone.0015077.

103. Blanchard, R.J., and Blanchard, D.C. (1969). Crouching as an index of fear. J Comp Physiol Psychol 67, 370–375. 10.1037/h0026779.

104. Grieves, R.M., Jedidi-Ayoub, S., Mishchanchuk, K., Liu, A., Renaudineau, S., and Jeffery, K.J. (2020). The place-cell representation of volumetric space in rats. Nat Commun 11, 789. 10.1038/s41467-020-14611-7.

