## Supplementary Figures 1-7 for "Disruption of hippocampal-prefrontal neural dynamics and risky decision-making in a mouse model of Alzheimer’s disease"

This pdf file includes 7 supplementary figures.

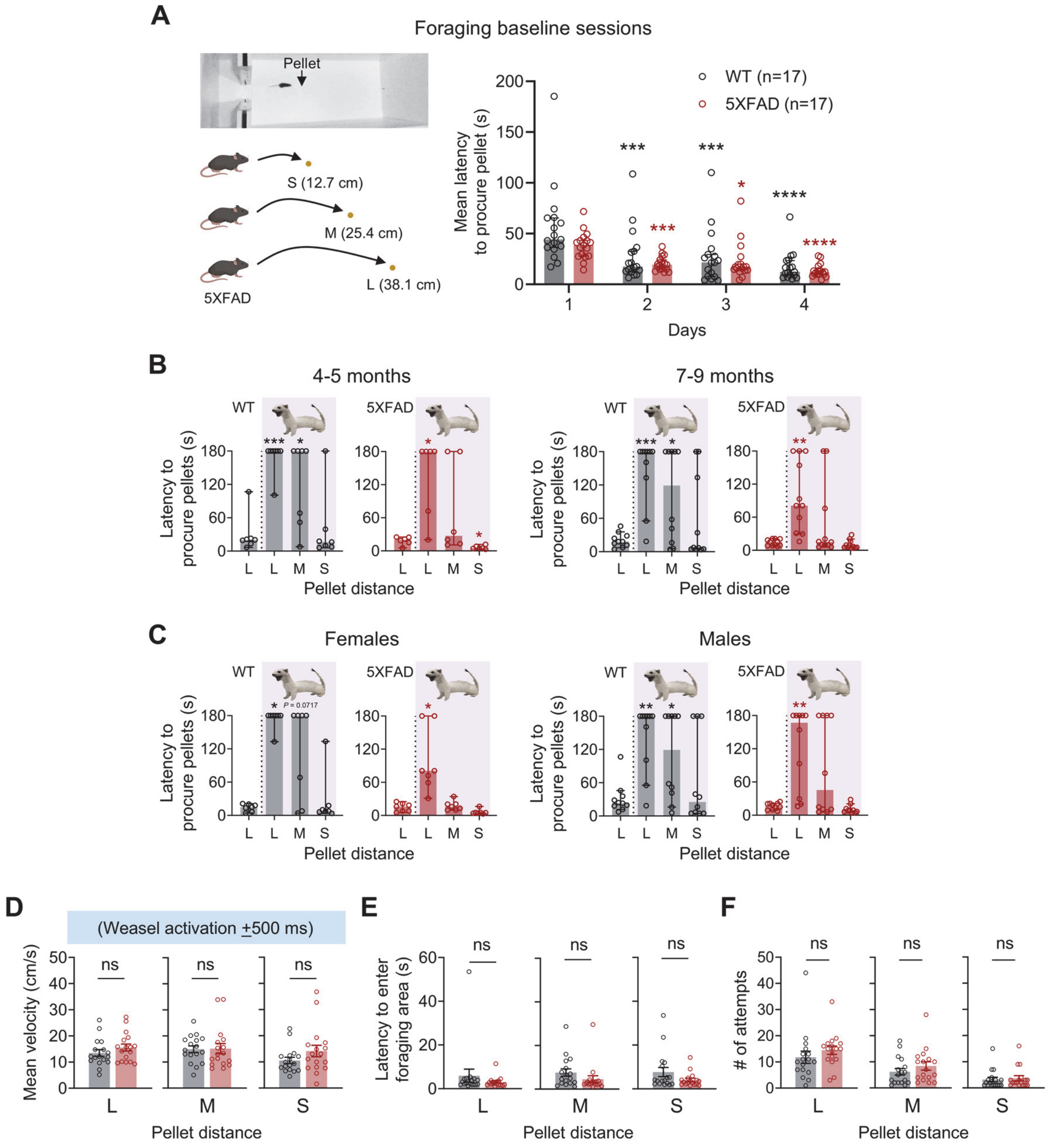


**FIGURE S1. Baseline foraging behavior and age/sex comparisons in risky decision making in 5XFAD mice.** (A) *Left*: Image of a mouse foraging during a baseline session, alongside the associated behavioral procedure. Animals underwent three trials where pellets were placed at increasing distances from the nest (12.7 cm, S; 25.4 cm, M; 38.1 cm, L), with 3 min allocated to procure each pellet. *Right*: Latency to procure pellets decreased over the baseline days for both WT and 5XFAD mice, with no significant differences between the groups. (WT: ***$P < 0.001$, ****$P < 0.0001$; 5XFAD: ***$P < 0.001$, ****$P < 0.0001$; compared to day 1) (B,C) Both 4-5-month-old and 7-9-month-old WT mice (B) exhibited longer latencies to procure the L and M pellets compared to their baseline levels, while 5XFAD mice showed no significant change in latency for M pellets, suggesting riskier decision-making in 5XFAD mice. This pattern persisted across both sexes in the WT and 5XFAD groups (C). (D-F) No significant group differences were observed in approach-escape speed across the three distance trials (around −500 and 500 ms from weasel activation; D), latency to enter the foraging area (E), and the number of pellet retrieval attempts (F) between 5XFAD and WT mice during predator trials. *$P < 0.05$, **$P < 0.01$, ***$P < 0.001$ compared to baseline latency.

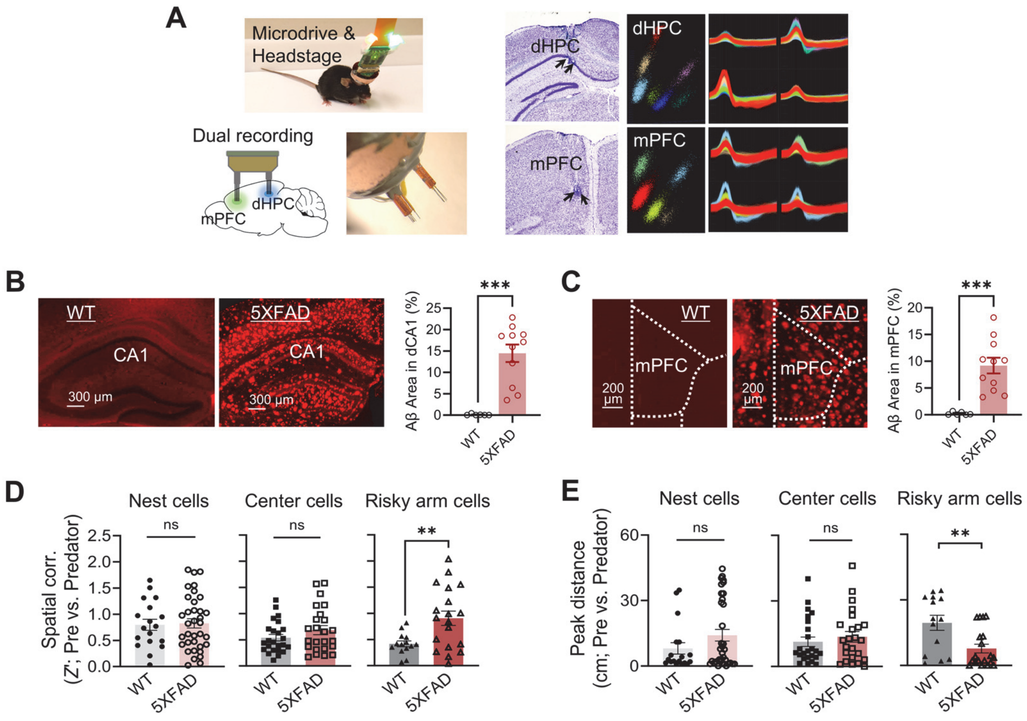


**FIGURE S2. dHPC-mPFC dual recording, Aβ deposits, and spatial analysis** (A) Photomicrographs of tetrodes implants and representative multiple single units recorded in the dHPC and mPFC (indicated by arrows). (B,C) Aβ accumulation in dCA1 (B) and mPFC (C) of 5XFAD and WT mice. (D) Session-by-session spatial correlation (Z') analysis revealed no significant group differences in nest and center cells, but significant differences were noted in risky arm cells. (E) Similar peak distances in nest and center cells across sessions, with significant differences in risky arm cells.

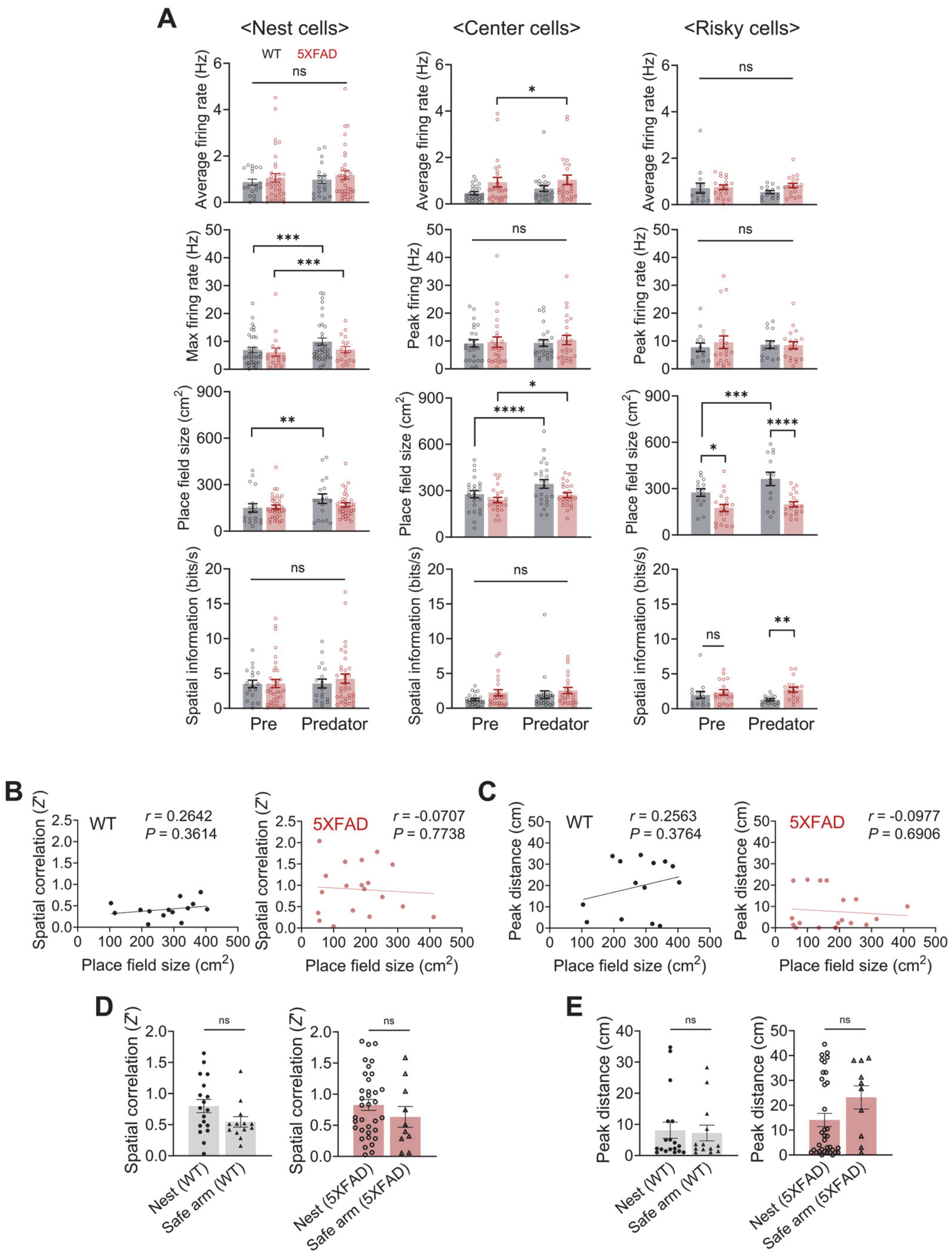


**FIGURE S3. Basic properties of dHPC place cells.** (A) Analysis of average firing rate, maximal firing rate, place field size, and spatial information for place cells in the nest, center, and risky arm from WT and 5XFAD mice. Average firing rate: No differences between groups and sessions. Maximal firing rate: Both WT and 5XFAD nest cells displayed an increased maximal firing rate during the predator session, while cells in the center and risky arm did not. Place field size: WT nest cells showed an increased place field size during the predator session compared to the pre-predator session. Both WT and 5XFAD center cells exhibited increased place field sizes in the presence of the predator. 5XFAD risky arm cells displayed consistently smaller place field sizes, yet both WT and 5XFAD risky arm cells demonstrated increases during the predator session. Spatial information: No significant differences were noted, except for higher spatial information in 5XFAD risky arm cells compared to WT during the predator session. (B) Place field size was not correlated with spatial correlation (Z') between pre-predator and predator sessions in risky arm cells of both WT and 5XFAD mice. (C) Place field size showed no correlation with peak distance between pre-predator and predator sessions in risky arm cells of both WT and 5XFAD mice. (D) Spatial correlation analysis revealed no significant differences between nest and safe arm cells in both WT and 5XFAD mice. (E) Peak distance comparisons also indicated no significant differences between cell types in both mouse groups.*$P$ < 0.05, **$P$ < 0.01, ***$P$ < 0.001, ****$P$ < 0.0001.

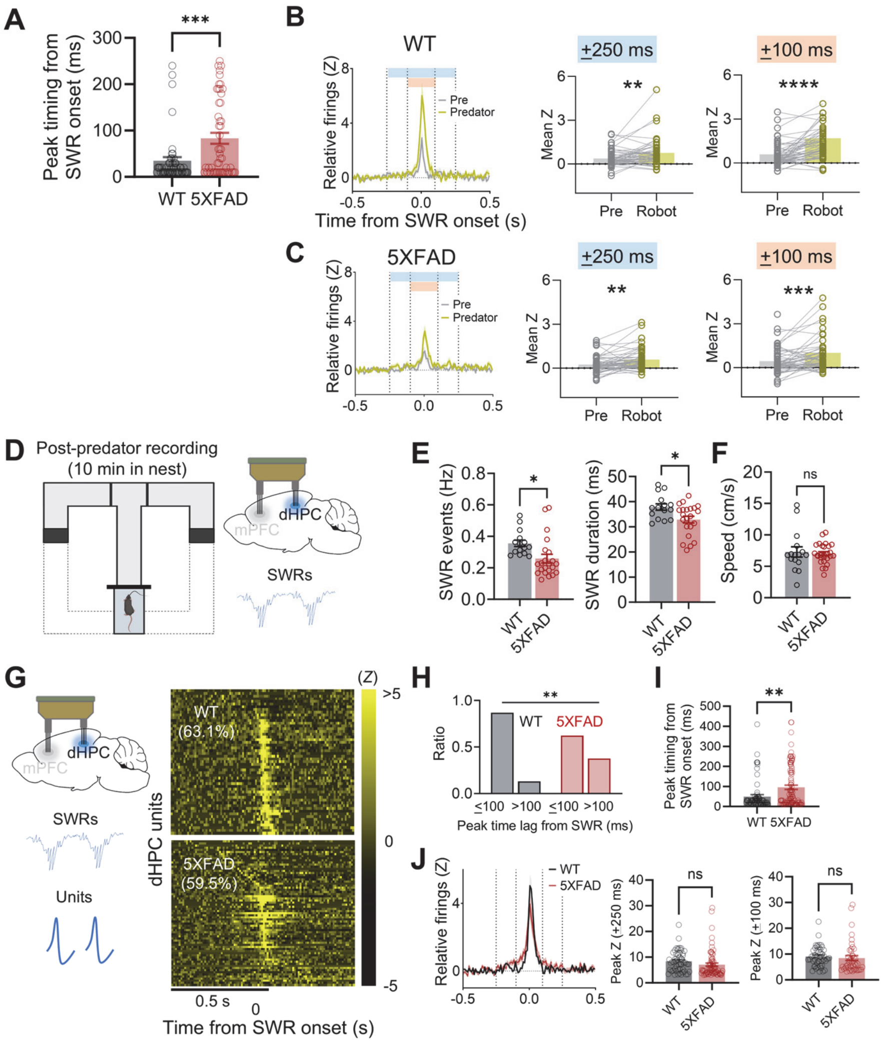

**FIGURE S4. SWR-related dHPC cell firing peak timing in WT and 5XFAD mice.** (A) 5XFAD dHPC cells fired farther from the SWR onset compared to WT dHPC cells. (B,C) Peri-SWR activity (mean *Z*-scores) of WT (B) and 5XFAD (C) dHPC neurons showing significant activity ($z > 3$) within the ±250 ms (blue) and ±100 ms (orange) periods around SWR onset during the predator session. Mean *Z*-scores were compared bewteen sessions. (D) Following the conditional predatory task, animals were confined to the nest for a 10-min post-predator encounter recording session. (E) The frequency (*Left*) and duration (*Right*) of SWRs significantly decreased during the post-predator encounter. (F) Overall speed did not differ between the WT and 5XFAD groups during the post-predator session. (G) dHPC SWR and unit activities were analyzed during the pre-predator and predator sessions, highlighting color-coded dHPC unit activity (significant Z scores) during peri-SWR epochs, aligned with SWR onset in each session. (H) Proportions of dHPC neurons exhibiting significant activity within the ±250 ms vs. ±100 ms periods differed between the WT and 5XFAD groups. (I) 5XFAD dHPC cells fired farther from the SWR onset compared to WT dHPC cells. (J) SWR-related dHPC cell activity (peak Z-scores within the ±250 ms but not ±100 ms periods) during the post-predator session did not differ between the groups. * $P < 0.05$, **$P < 0.01$, ***$P < 0.001$, ****$P < 0.0001$.

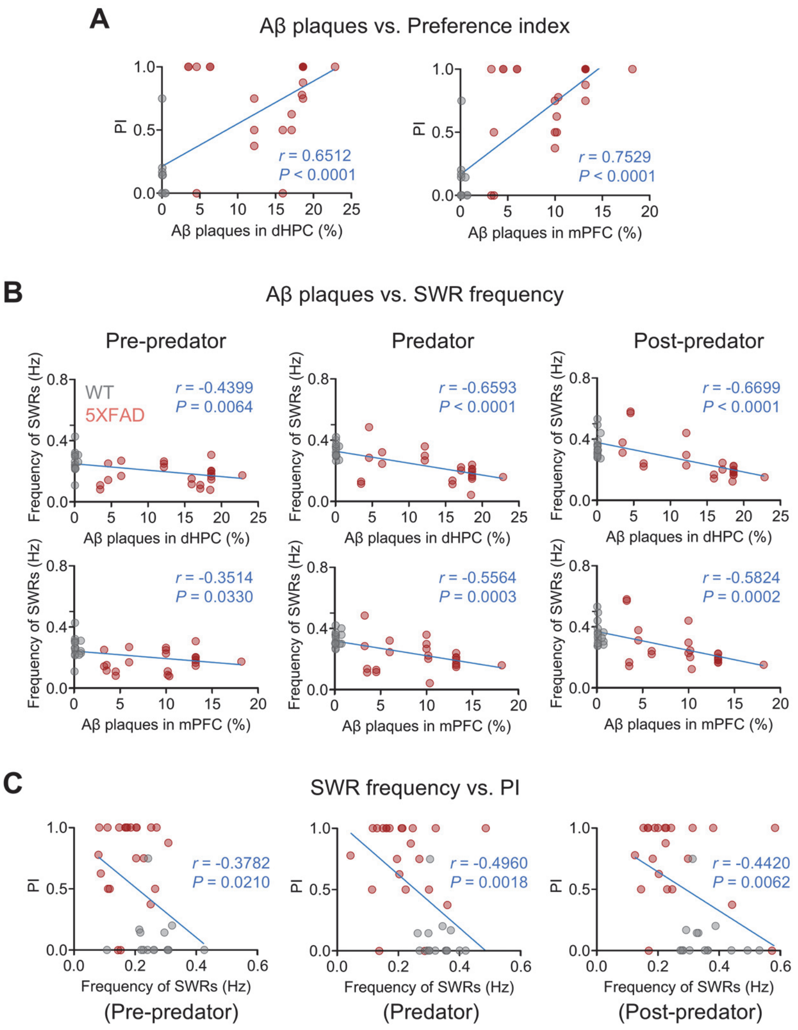


**FIGURE S5. The relationship between amyloid load, SWR activity, and behavioral flexibility.** (A) Significant correlations were found between Aβ plaque levels in the dHPC and mPFC and the behavioral index (PI) value. (B) Aβ plaque deposition levels in both dHPC and mPFC showed significant correlations with SWR frequency across all sessions. (C) SWR frequency in all sessions showed a significant correlation with PI values.

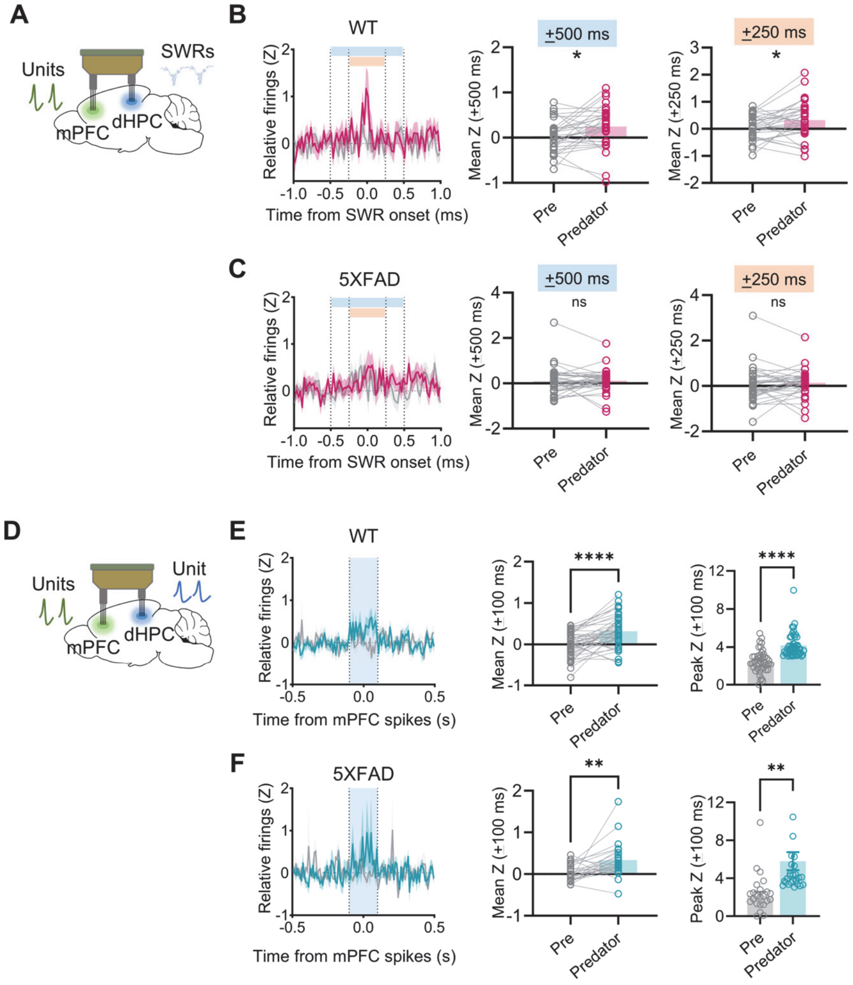


**FIGURE S6. Alterations in dHPC-mPFC dynamics during risky decision-making in 5XFAD mice.** (A) dHPC SWR and mPFC unit activities were analyzed during pre-predator and predator sessions, exploring how dynamic interactions are altered in risky contexts. (B) Peri-SWR activity (mean Z-scores) of WT mPFC neurons exhibiting significant activity ($z > 3$) within ±500 ms (blue) and ±250 ms (orange) around SWR onset during the predator session. Mean *Z*-scores were compared between sessions to assess changes. (C) Similar peri-SWR activity (mean *Z*-scores) analysis for 5XFAD mPFC neurons, with comparisons of mean Z-scores during ±500 ms and ±250 ms between sessions to identify differences in response patterns. (D) dHPC and mPFC unit activities were analyzed during the pre-predator and predator sessions to understand inter-regional neural interactions. (E,F) Averaged dHPC-mPFC CCs from WT (E) and 5XFAD (F) mice showing significant peaks (between −100 and 100 ms; blue area) during pre-turn epochs in the presence of the predator (*Left*). Mean (*Middle*) and peak (*Right*) *Z*-scores within the +100 ms period during both pre-predator and predator sessions in WT and 5XFAD mice. Shaded areas indicate SEM. *$P$ < 0.05, **$P$ < 0.01, ****$P$ < 0.0001.

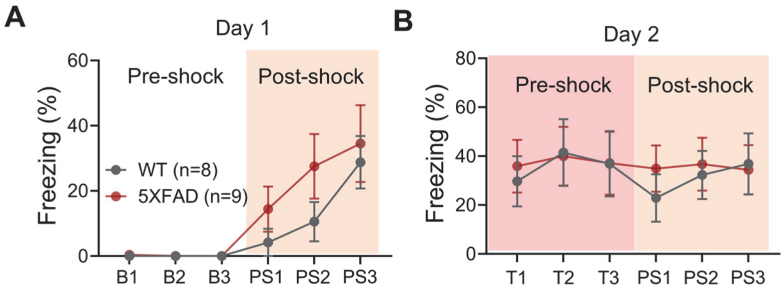


**FIGURE S7. Contextual fear conditioning in 5XFAD mice.** (A,B) During days 1 (A) and 2 (B), both 5XFAD and WT mice exhibited similar levels of freezing in the pre-shock (3 min) and post-shock (3 min) phases, indicating comparable fear conditioning responses across both groups.
